# Eomes directs the formation of spatially and functionally diverse extra-embryonic hematovascular tissues

**DOI:** 10.1101/2024.08.13.607790

**Authors:** Bart Theeuwes, Luke TG Harland, Alexandra Bisia, Ita Costello, Mai-Linh Ton, Tim Lohoff, Stephen J Clark, Ricard Argelaguet, Nicola K Wilson, Wolf Reik, Elizabeth Bikoff, Elizabeth J Robertson, Berthold Gottgens

## Abstract

During mouse gastrulation, extraembryonic mesoderm (ExEM) contributes to the extraembryonic yolk sac (YS) and allantois, both of which are essential for successful gestation. Although the genetic networks coordinating intra-embryonic mesodermal subtype specification are well-studied, the mechanisms driving ExEM diversification are poorly understood. Here, we reveal that embryoid body *in vitro* differentiation generates two distinct lineages of mesodermal cells matching YS and allantois respectively. Combining *in vitro* models with *in vivo* chimeric embryo analysis, we discover that Eomesodermin (Eomes) regulates the formation of a subset of YS-fated ExEM but is dispensable for allantois formation. Furthermore, simultaneous disruption of Eomes and T impedes the specification of any YS or allantois mesoderm, indicating compensatory roles for T during allantois formation when Eomes is disrupted. Our study highlights previously unrecognized functional and mechanistic diversity in ExEM diversification and endothelial development and introduces a tractable EB model to dissect the signaling pathways and transcriptional networks driving the formation of key extraembryonic tissues.

## Introduction

At the onset of murine gastrulation, extraembryonic mesodermal (ExEM) cells originating from the most posterior region of the primitive streak (PS) migrate proximally and form hematovascular and mesenchymal cell types of both the visceral yolk sac (YS) and allantois (Fig 1A). YS mesoderm gives rise to an extensive endothelial vascular plexus that regulates nutrient exchange and functions as the initial site of hematopoiesis during embryogenesis (Dzierzak and Bigas, 2018). The allantoic bud arises from a progenitor pool at the proximal posterior of the PS, that proliferates, extends and fuses with the chorionic plate to form the chorioallantoic placenta by E8.5 (Downs, 2022; Inman and Downs, 2007). At later developmental stages, allantois mesoderm forms the umbilical cord and fetal placental vasculature (Arora and Papaioannou, 2012), but its potential to generate *de novo* hematopoietic progenitors is less well understood. Despite its crucial role in vasculature formation and establishing the fetal-maternal blood supply, the allantois has often been overlooked in embryological studies (Downs, 2022; Inman and Downs, 2007) and during generation of hematovascular progenitors in embryonic stem cell (ESC) cultures (Ditadi et al., 2017). Consequently, models of the allantois are underdeveloped, and the molecular pathways governing ExEM diversification into distinct extraembryonic lineages remain poorly understood.

**Figure 1:**
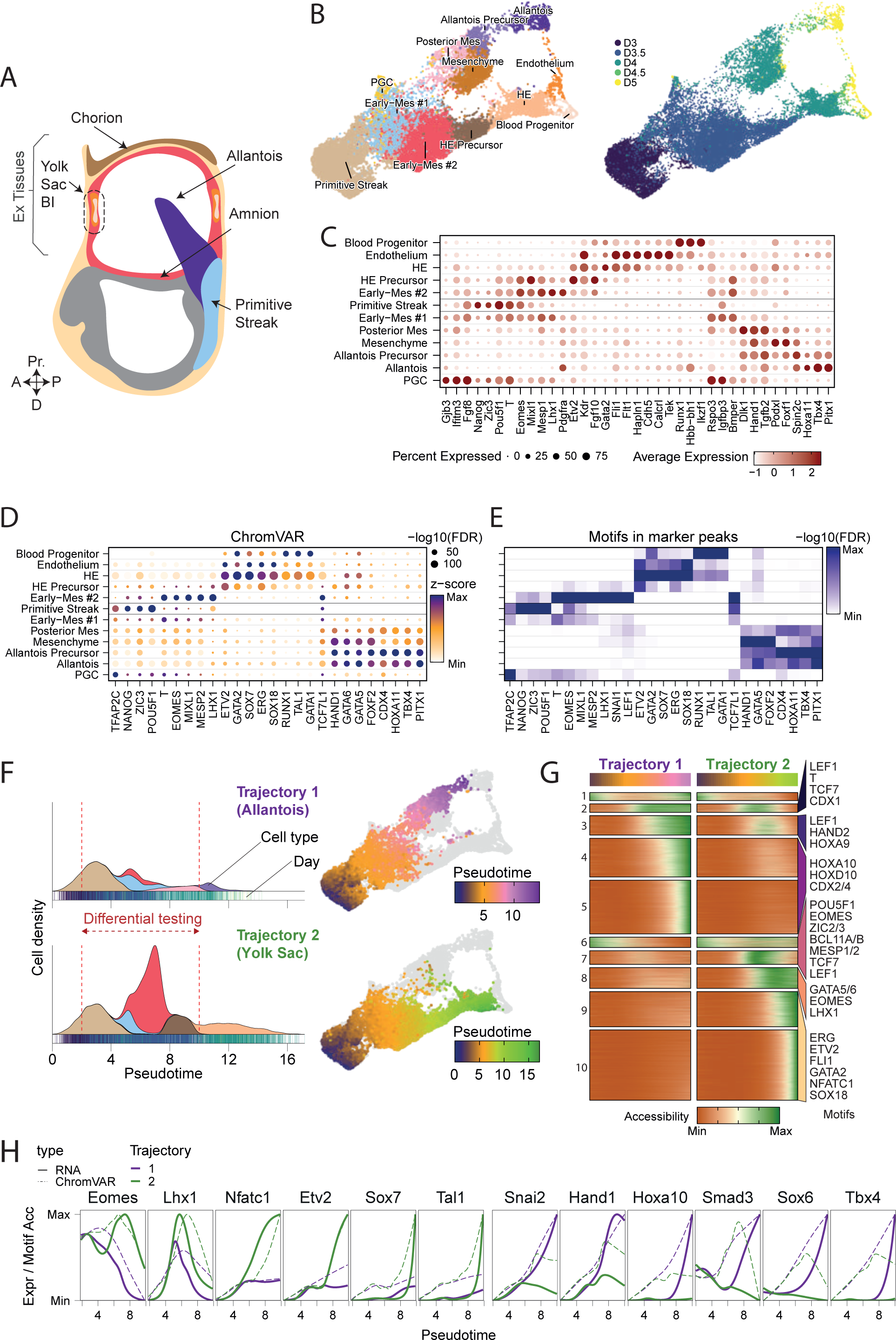
Multiomic characterization of embryoid bodies reveals yolk-sac and allantois lineage differentiation. A) Schematic of an E8.25 mouse gastrula. Ex = extraembryonic, Pr = proximal, A = anterior, P = posterior and D = distal, BI = blood islands. B) Uniform Manifold Approximation and Projection (UMAP) of shared scRNA-seq and scATAC-seq, colored by cell type annotation (left) and day (right). D = day. C-E) Molecular characterization of EB cell types, displaying marker gene expression (C), ChromVAR scores (D) and motif enrichment in ATAC marker peaks (E). F) Left-side, cell type densities displayed along pseudo times for Allantois Trajectory #1 (top) and YS Trajectory #2 (bottom). Right-side, pseudo times for Trajectory #1 (top) and Trajectory #2 (bottom) overlaid on UMAPs. G) Heatmap showing normalized chromatin accessibility for peaks along pseudo time for both trajectories, clustered by peak accessibility patterns. Enriched motifs for cluster(s) are displayed on the right. Differential testing was performed on pseudotime range indicated in F. H) Gene expression (solid lines) and chromVAR motif accessibility (dashed lines) for allantois (purple – trajectory 1) and YS (green – trajectory 2) trajectories plotted along pseudotime.

T-box transcription factors (TF) Eomesodermin (Eomes) and Brachyury (T) are dynamically expressed in the epiblast, primitive streak, and ExEM precursors at the onset of murine gastrulation (Arnold et al., 2008; Ciruna and Rossant, 1999; Costello et al., 2011; Harland et al., 2021; Herrmann et al., 1990; Rivera-Pérez and Magnuson, 2005). Disrupting the functionality of Eomes or T in mouse embryos and ESC cultures has revealed their essential and partially redundant roles during gastrulation (Arnold et al., 2008; Costello et al., 2011; Harland et al., 2021; Schüle et al., 2023; Tosic et al., 2019) T is dispensable for the initial specification of anterior intraembryonic mesoderm and ExEM but is required for sustained posterior mesoderm development and axial elongation (Guibentif et al., 2021; Rashbass et al., 1991; Wilson et al., 1993), while Eomes is crucial for specifying cardiac mesoderm and the definitive endoderm lineage (Arnold et al., 2008; Costello et al., 2011). Using an *in vitro* model of hematovascular formation, we recently demonstrated that Eomes is necessary for YS hematopoiesis and hemogenic endothelial formation but dispensable for non-hemogenic endothelium (Harland et al., 2021). Interestingly, *in vivo* fate mapping experiments demonstrate that Eomes+ cells contribute to the allantois (Harland et al., 2021), however, the role of Eomes during allantois formation has not been investigated. More broadly, the underlying Eomes-dependent mechanisms that coordinate formation of early hematovascular lineages remain poorly understood and a cell-autonomous role for Eomes during blood and endothelial formation *in vivo* has yet to be determined.

Here we utilized 10x Genomic’s multiome to create a detailed, time-resolved single-cell atlas of gene expression (RNA) and chromatin accessibility (ATAC) during murine hematovascular embryoid body (EB) differentiation. This approach revealed robust production of both YS and allantois cell types. Using a combination of *in vitro* and *in vivo* assays, we demonstrate that Eomes is essential for the formation of YS-fated ExEM, but surprisingly is not required for specification of the allantois lineage and allantois endothelium. Eomes dependence thus clearly defines the formation of distinct extraembryonic hematovascular tissues. The previously described block in YS hematopoiesis, caused by Eomes loss-of-function, reflects an earlier loss in YS-fated ExEM. Moreover, we demonstrate that simultaneous disruption of Eomes and T entirely blocks the specification of all YS and allantois mesoderm. Taken together, these findings reveal new insights into the mechanisms that allow TFs to coordinate ExEM diversification. Additionally, the ability to readily generate allantoic mesoderm during ESC differentiation *in vitro* has important implications for future studies of developmental biology and regenerative medicine.

## Results

### Efficient generation of extra-embryonic mesoderm sub-types in vitro

We recently described a mouse ESC differentiation protocol that robustly generates extra-embryonic populations, including YS-like haematopoietic progenitors, via the formation of EBs (Harland et al., 2021). However, a comprehensive and unbiased analysis exploring how this *in vitro* system recapitulates *in vivo* embryonic development is missing. Here, using multiomics, we characterized the transcriptional/chromatin accessibility profiles of over 19,000 single cells, isolated at 12-hour intervals from days 3 to 5 of EB differentiation (Fig 1B, SFig 1A-D). We annotated 12 cell types using marker genes and motif accessibility patterns as well as label transfer from *in vivo* reference atlases (Argelaguet et al., 2022; Imaz-Rosshandler et al., 2024; Pijuan-Sala et al., 2019) (Fig 1B-E). Additionally, we used in-silico ChIP methodology to predict TF occupancy (Fig 1D, E; Argelaguet et al., 2022, see Methods).

At day 3, EBs were primarily composed of PS cells expressing Nanog, Zic3, and Pou5f1 (Fig 1B,C). Twelve hours later, two early mesoderm (Early-Mes) cell populations emerged, which we labelled Early-Mes #1 and Early-Mes #2. The transcriptomes of both populations were highly similar. However, Early-Mes #1 expressed higher levels of Rspo3, and Igfbp3 while Early-Mes #2 expressed higher levels of Eomes, Mixl1, Mesp1, Lhx1, and Pdgfra (Fig 1C). Day 3.5 EBs contained hematoendothelial (HE) precursors expressing Early-Mes #2 markers as well as Kdr, Etv2, and Fgf10 (Fig 1C).

Beginning at day 4, we detected HE cells expressing Etv2, Kdr, Fli1, and Gata2 with regions of increased chromatin accessibility enriched for ETV2, GATA2, and ERG motifs (Fig 1B-E). During early gastrulation, RNA expression of Tbx4 and Hoxa10/11/13 are exclusively localized to the allantois bud (Naiche et al., 2011; Scotti et al., 2015). Our day 4 EBs contained allantois-associated cell types, including a posterior mesodermal subset (Dlk1, Hand1), allantois precursors (Tbx4, Pitx1, and TBX4/PITX1/HOXA11 motifs), as well as mesenchymal cells (Hand1, Podxl, Foxf1, and FOXF/GATA5/6 motifs) (Fig 1B-E).

By day 4.5/5, EBs contained blood progenitors (Runx1, Hbb-bh1, Ikzf1, and RUNX1/GATA1/TAL1 motifs), endothelial cells (Cdh5, Tek, Fli1, and ETV2/SOX/ERG motifs), mesenchyme, and an allantois-like cell population (Tbx4/Pitx1 and CDX/HOX/PITX1 motifs) (Fig 1B-E). We detected a few PGC-like cells, expressing Gjb3 and Ifitm3 and enriched for TFAP2C motifs, from day 3.5 onwards (Fig 1B-E). Notably, the chromatin landscapes of EB cell types closely mirrored their *in vivo* counterparts (SFig 2A). Altogether, these findings demonstrate our EB differentiation system effectively recapitulates the formation of ExEM populations that form hematopoietic, endothelial, and mesenchymal cell types arising in both the YS and allantois.

### Multiome analysis identifies two distinct trajectories guiding the formation of the yolk-sac and allantois lineages

Next, we performed trajectory inference and identified two major trajectories (Fig 1F). Trajectory #1 generates allantois-like cells from posterior mesodermal precursors arising from Early-Mes #1, whereas Trajectory #2 resembles YS formation including downstream blood progenitors and endothelial cells generated from HE precursors arising from Kdr+ Early-Mes #2. Notably, Early-Mes #1 is included in both trajectories prior to an early (D3.5) lineage bifurcation (Fig 1B,F). The PS/Early-Mes #1 are therefore likely common precursors for both lineages, consistent with *in vivo* transplantation studies showing continuous generation of YS- and allantois-fated mesoderm from the posterior PS during gastrulation (Kinder et al., 1999).

Using TradeSeq, we uncovered gene sets and chromatin peaks that display distinctive patterns of expression/accessibility across the pseudotimes for both trajectories (Fig 1G,H, SFig 2B,C). Eomes, Lhx1, and Etv2 increased in expression and motif accessibility at the onset of the YS trajectory, whereas Snai2 and Hand1 were upregulated early in the allantois trajectory (Fig 1G,H). At later pseudotimes, Sry-box TFs, Sox7 and Sox6, exhibited specificity for the YS and allantois trajectories, respectively (Fig 1H). Signalling-related genes were also differentially expressed. Early in the YS trajectory, the ligand Vegfc and its receptor Kdr were upregulated (SFig 2C), consistent with lineage tracing experiments tracking Kdr+ mesoderm to YS blood islands (Lugus et al., 2009). Wnt and Fgf receptors/ligands (Wnt: Fzd7, Frzb, Wnt5a, and Lgr4; Fgf: Fgf10/15/18) as well as Tgf-β/BMP pathway elements including receptors (Tdgf1 and Tgfbr3), inhibitors (Lefty2 and Bambi), ligands (Bmp2, Bmp7, and Tgfb2), and downstream effectors (Smad9) also exhibit distinctive expression patterns (SFig 2C). Overall, our analyses reveal dynamic and distinctive expression patterns of inductive cues and transcription factors, which potentially direct cell fate specification at various stages of YS and allantois differentiation.

### Transient Eomes expression in ExEM progenitors is required for the YS lineage but dispensable for allantois cell populations

Using Eomes knock-out (KO) ESC, we previously identified an essential role for Eomes in YS-like hematopoiesis and bulk chromatin changes of D4 KDR+ cells suggested Eomes primes early mesoderm with hemogenic competence (Harland et al., 2021). The trajectory analyses presented above suggest Eomes acts at the onset (D3-D3.5) of ExEM diversification, when bifurcation towards the YS versus allantois lineage occurs (Fig 1H). To further investigate Eomes requirements during this early time-window, we employed flow cytometry, functional assays, and multiomics using a Runx1^Venus^ Eomes^mCherry-degron^ ESC line (Fig 2A) (Bisia et al., 2023). This line features a degron tag attached to the C-terminus of Eomes, which enables rapid EOMES depletion via adding the small molecule dTAG13 to culture media (Bisia et al., 2023). Additionally, the Venus reporter in this line monitors Runx1 expression, which marks nascent hematopoietic progenitors (Yzaguirre et al., 2017).

**Figure 2:**
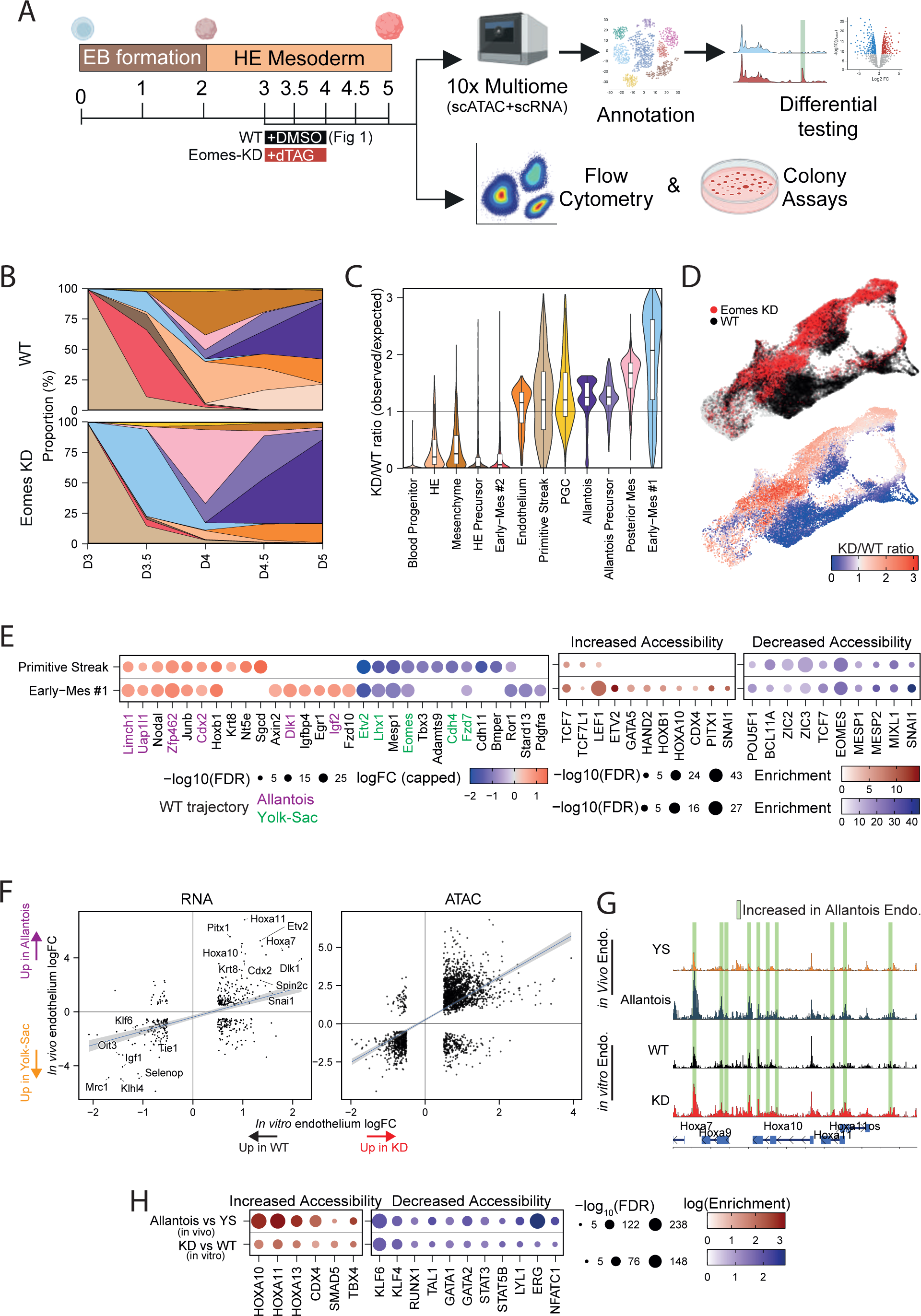
Allantois cell types predominate downstream of Eomes protein knockdown in EB cultures due to a disruption in early mesoderm formation. A) Experimental design: 10x Genomics multiome sequencing, flow cytometry, and colony assays were conducted on WT and Eomes-KD EB cultures. WT cultures received DMSO and Eomes-KD cultures received dTAG13 from D3 to D4. B) Cell type proportions per timepoint for WT (top) and Eomes KD cultures (bottom). At D3 WT proportions are displayed, as Eomes KD was induced from D3 onwards. C) Ratio of Eomes-KD to WT cells, calculated using the 100 nearest neighbors for each cell, colored by cell type. Values above one indicates enrichment of Eomes-KD cells, while values below one indicates depletion. D) UMAPs colored by EB culture condition (top) and Eomes KD to WT ratio as shown in C (bottom). E) Left panel shows differential gene expression between Eomes-KD and WT early cell types. Colored gene names indicate allantois (purple) and YS (green) marker genes from Fig 1. Middle/right panels display motif enrichment for peaks with increased (middle) or decreased (right) chromatin accessibility. F) Differentially expressed genes (left) and differentially accessible peaks (right) for Eome*s-* KD versus WT endothelium (x-axis) compared to *in vivo* allantois versus YS endothelium (y-axis). G) Chromatin accessibility at the Hoxa locus for *in vivo* YS and allantois endothelium (endo.) and *in vitro* WT and Eomes-KD endothelium. Green bars highlight peaks with increased chromatin accessibility in allantois endothelium. H) Motif enrichment for peaks with increased (left) or decreased (right) accessibility between endothelial cells of the in vivo allantois and YS from Argelaguet et al., 2022 (top row) and the in vitro Eomes-KD and WT cells (bottom row).

At day 3 of EB differentiation, flow cytometry revealed ∼70% of cells expressed EOMES protein prior to upregulation of mesodermal markers KDR and PDGFRA (SFig 3A,B). The proportion of EOMES-expressing cells dropped substantially by day 4 (∼15%) and by day 4.5 EOMES was no longer detectable (SFig 3A). Addition of dTAG13 from day 3 to 4, hereafter referred to as Eomes knockdown (Eomes-KD), rapidly abolished EOMES protein, resulting in Eomes depletion and notably blocked formation of a day 3.5 KDR^+^PDGFRA^+^ population (SFig 3A,B). Additionally, in line with previous Eomes-KO experiments (Harland et al., 2021), Eomes-KD abrogated formation of Runx1-Venus+/CD41+ blood progenitors and primitive erythrocytes, but failed to block generation of CDH5+ endothelial cells (SFig 3C,D). Kinetics of RNA versus protein expression for these markers were highly similar during WT EB differentiation (SFig 3A,B,D,E).

To further investigate the molecular consequences of Eomes-KD, we conducted multiomics profiling of Eomes-KD EBs at 12-hour intervals from days 3.5 to and compared these profiles to equivalent DMSO-treated WT EBs shown in Fig 1 (Fig 2A). Strikingly, Eomes-KD cells lack the ability to contribute to cell types represented in the YS trajectory (Fig 2B-D). By day 3.5, Eomes-KD cultures generated very few, if any, Early-Mes #2 and HE precursors, and at later stages failed to form HE and blood progenitors. By contrast, cell types associated with the allantois trajectory, including Early-Mes #1, posterior mesoderm, allantois precursors, and the Tbx4+ allantois cell population, were abundantly generated (Fig 2B-D). These results demonstrate that transient Eomes expression governs the formation of an early subset of YS-fated ExEM and its downstream progeny but is dispensable for allantois formation.

To further explore Eomes’ role during the initial step of ExEM diversification, we examined Eomes-dependent molecular changes in PS and Early-Mes #1 cell types, common precursors for both trajectories (Fig 2E). Notably, Eomes disruption caused immediate downregulation of Etv2, a key regulator of YS hematovascular formation (Koyano-Nakagawa and Garry, 2017; Koyano-Nakagawa et al., 2012; Liu et al., 2015; Rasmussen et al., 2011), as well as Eomes target genes Lhx1 and Mesp1 (Costello et al., 2011; Nowotschin et al., 2013) (Fig 2E). Additionally, motifs for EOMES, mesoderm-related TFs (MESP1/2, MIXL1, SNAI1) and pluripotency factors (ZIC2/3, POU5F1) were enriched in regions with decreased chromatin accessibility (Fig 2E). Conversely, Eomes-KD led to premature upregulation of TFs and signaling molecules (Cdx2, Nodal, Dlk1, and Igf2) (Fig 2E, SFig 3F) as well as increased chromatin accessibility enriched for motifs (HOX, CDX4 and PITX1) linked to later stages of the allantois trajectory (Fig 1D,E,G). Interestingly, Eomes expression was also reduced, implying an autoregulatory mechanism likely drives Eomes expression during YS formation (Fig 2E). Thus, Eomes regulates chromatin and transcriptional biases during ExEM diversification that pattern allantois versus YS fates in an early posterior PS population.

### Specification of YS but not allantoic endothelium is Eomes dependent

Both the YS and allantois are major sites of *de novo* vasculogenesis during embryogenesis and reference atlases have revealed molecular signatures that distinguish endothelium arising in these tissues (Imaz-Rosshandler et al., 2024). Eomes-KD and WT EBs produced similar proportions of endothelial cells; however, Eomes-KD impedes YS but not allantois differentiation (Fig 2C). To investigate endothelial diversity in EBs, we compared the molecular differences between *in vivo* allantois and YS endothelium to those between *in vitro* Eomes-KD and WT endothelium (Fig 2F).

Eomes-KD endothelial cells exhibited elevated levels of allantois endothelial marker genes (e.g., Dlk1 and Hoxa7/10/11) and chromatin peaks (Fig 2F). Furthermore, sites of increased chromatin accessibility in Eomes-KD and allantois endothelial cells were enriched for the same motifs (HOX, CDX4, and TBX4) (Fig 2H). Conversely, Eomes-KD endothelium showed reduced expression of YS endothelial marker genes (e.g. Mrc1, Klf6, Oit3, Tie1), and sites of decreased chromatin accessibility in Eomes-KD endothelium were enriched for YS endothelial motifs (KLF6, RUNX1, STAT5B, TAL1, and LYL1) (Fig 2F, H). Furthermore, comparing specific genomic regions (e.g. HOXA locus) highlights the allantoic endothelial signature of Eomes-KD endothelium (Fig 2G, SFig 3G). Interestingly, allantois marker peaks were accessible in WT endothelial cells, albeit at lower levels compared to Eomes-KD endothelium (Fig 2G and SFig 3G). Altogether, these analyses suggest that WT EB cultures contain both YS and allantois-like endothelial cells. In contrast, Eomes-KD EBs selectively generate allantois-like endothelium, and lack endothelium with a YS molecular signature, likely due to the earlier block in YS ExEM differentiation described above. Interestingly, Eomes-KD endothelium exhibited increased Etv2 expression (Fig 2F), indicating that Eomes is not required for Etv2 induction in the allantois lineage during endothelial formation.

### Chimeric embryo analyses validate critical functions of Eomes for yolk sac but not allantois formation

To test the utility of our EB model and validate these findings *in vivo*, we conducted single-cell RNA sequencing (scRNA-seq) of chimeric embryos generated by injecting Eomes loss-of-function ESC expressing TdTomato into WT blastocysts (Fig 3A, SFig 4A). At E8.25-E8.5, chimeric embryos were dissociated, WT host cells and TdTomato+ Eomes-KO cells were sorted using FACS and analysed by 10X Genomics scRNA-seq (Fig 3A). The transcriptional profiles of WT and Eomes-KO cells from chimeric embryos were aligned with a gastrulation reference atlas (Pijuan-Sala et al., 2019), enabling cell type annotation and identification of cellular and molecular disruptions caused by Eomes loss-of-function (Fig 3B, SFig 4B).

**Figure 3:**
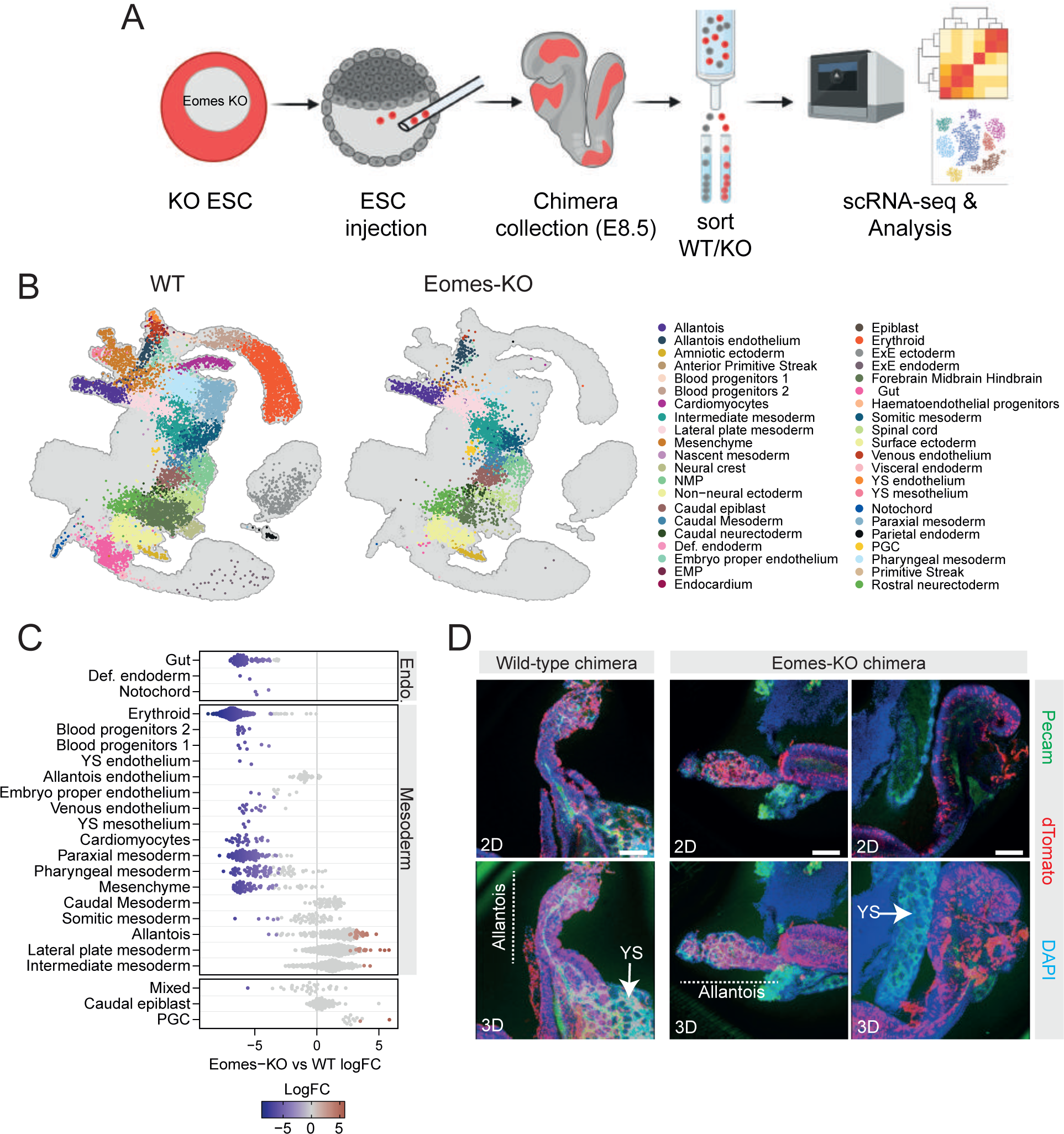
Eomes is essential for YS but dispensable for allantois development *in vivo*. A) Experimental design for Eomes-KO/WT chimaera-seq. Created with BioRender.com. B) Projection of chimera-seq WT (left) and Eomes-KO (right) cells onto the gastrulation atlas UMAP from Pijuan-Sala et al., 2019, colored by cell type annotation. C) Neighborhood-level differential abundance testing between Eomes-KO and WT cells. Negative logFC values indicate depletion of KO cells, while positive logFC values indicate enrichment, neighborhoods without statistical significance are colored grey. D) Confocal imaging of chimeras generated by injecting WT (left) or Eomes-KO (right) tdTomato+ ESCs into WT blastocysts. Samples are stained for DAPI (blue), injected tdTomato+ cells (red), and Pecam-1 (green). YS and allantois structures are indicated.

WT cells successfully mapped to the ectodermal, mesodermal, and endodermal cell types typically found in E8.25 embryos, as well as non-epiblast-derived lineages such as extraembryonic endoderm and ectoderm (Fig 3B). In contrast, Eomes-KO cells fail to form non-epiblast-derived lineages, reflecting the restricted potency of ESC (Fig 3B). To investigate statistical differences in the contribution of WT versus Eomes-KO cells to various cellular populations, we performed neighborhood-level differential abundance testing (Fig 3C and SFig 4C) (Dann et al., 2021). Consistent with previous fate mapping and functional studies (Arnold et al., 2008; Costello et al., 2011; Harland et al., 2021) Eomes-KO cells were deficient in generating gut tube and anterior mesodermal derivatives, such as cranial and pharyngeal mesoderm and cardiomyocytes (Fig 3B,C). However Eomes-KO cells efficiently contribute to posterior mesodermal derivatives, including caudal and somitic mesoderm, although formation of paraxial mesoderm was impaired (Fig 3B,C).

Consistent with our *in vitro* results, Eomes-KO cells failed to form YS endothelium and blood progenitors but were able to generate allantois and allantois endothelial cells (Fig 3B,C).

Furthermore, the transcriptional profiles of Eomes-KO versus WT allantois endothelium were highly concordant (SFig 4D). Numerous genes, however, were differentially expressed in allantois and lateral plate mesoderm cell populations (SFig 4D, E). Finally, confocal imaging of Eomes-KO chimeric embryos corroborated our scRNA-seq analyses, revealing a complete absence of tdTomato+ Eomes-KO cells in the YS but a significant contribution to the allantois, including Pecam1+ allantois endothelium (Fig 3D).

### Eomes and T play distinct and compensatory roles during ExEM formation

During gastrulation T expression is detected shortly after Eomes in the proximal posterior epiblast, nascent mesoderm along the length of the primitive streak as well as in the cell population at the base of the forming allantoic bud (Fig 4A) (Herrmann et al., 1990; Inman and Downs, 2007; Rivera-Pérez and Magnuson, 2005). We previously conducted scRNA-seq on T-KO chimeric embryos at E8.5 (Guibentif et al., 2021), which reveals no evident disruption of YS blood or endothelial cells or allantois cell types, including allantois endothelium (Fig 4B). These findings are consistent with earlier studies that show T-deficient ESC can contribute normally to the developing allantois region in chimeric embryos at E8.25 (Rashbass et al., 1991). Chorio-allantois fusion, which occurs at slightly later developmental stages is impaired in highly chimeric T-deficient allantoises, mimicking T-null embryos (Inman and Downs, 2006; Wilson et al., 1993).

**Figure 4:**
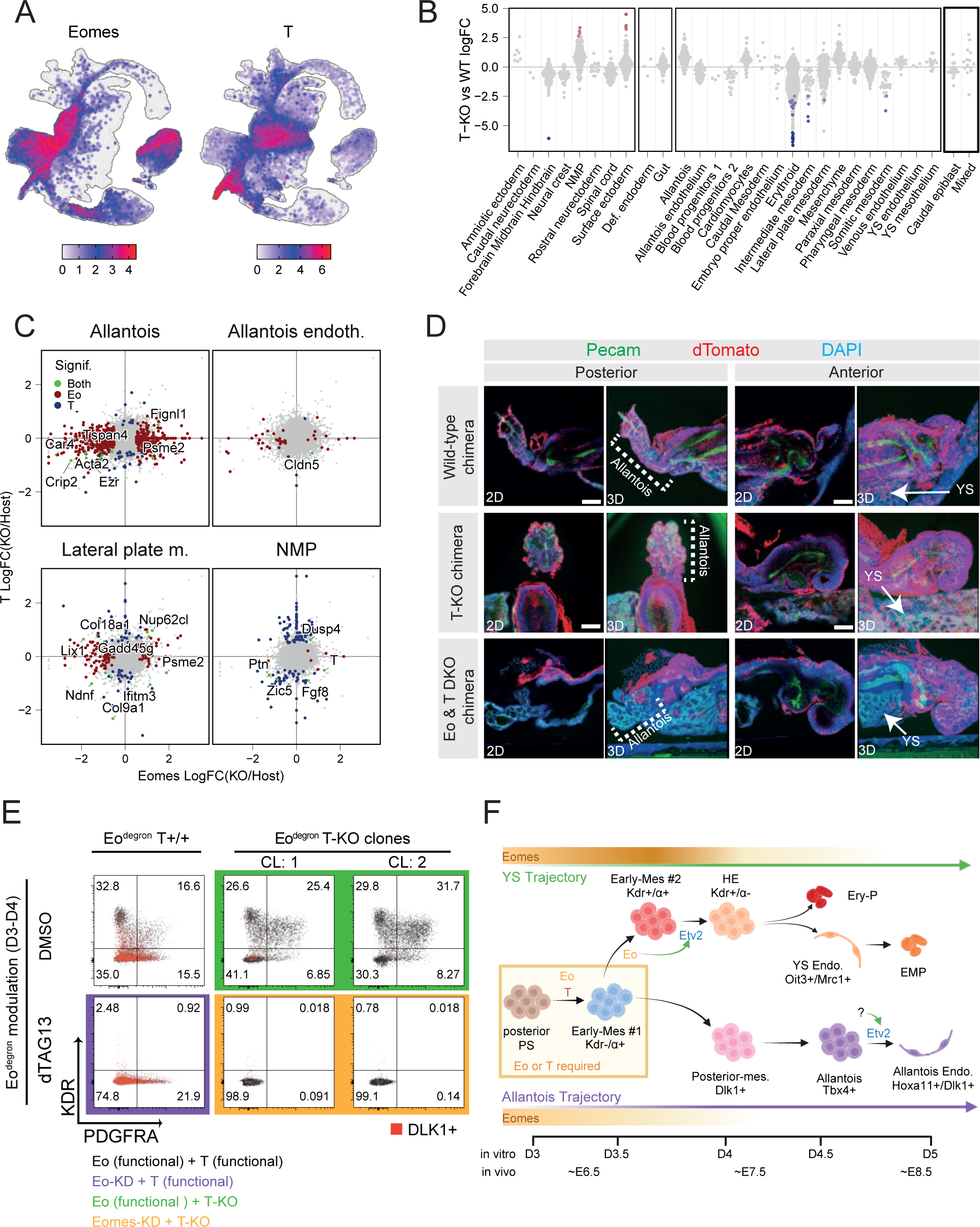
Eomes and T have distinct and compensatory roles during ExEM formation. A) Eomes and T RNA expression patterns displayed on the gastrulation atlas UMAP which covers E6.5-E8.5 of mouse development (Pijuan-Sala et al., 2019). B) Neighborhood-level differential abundance testing between T-KO and WT cells. Negative logFC values indicate depletion of KO cells, while positive logFC values indicate enrichment, neighborhoods without statistical significance are colored grey. C) Comparison of differentially expressed genes between injected KO and host WT cells for Eomes (x-axis) and T (y-axis) chimera-seq experiments, within specific cell types. Highlighted genes are significant in either T, Eomes, or both KO conditions. D) Confocal imaging of chimeras generated by injecting WT (row 1), T-KO (row 2), Eomes/T double KO (row 3) TdTomato+ ESC into a WT host blastocyst. Samples are stained for DAPI (blue), injected cells (red), and Pecam-1 (green). YS and allantois structures are indicated. E) Flow cytometric analyses of PDGFRA and KDR expression for Eomes^WT^/T^WT^, Eomes^KD^/T^WT^, Eomes^WT^/T^KO^, Eomes^KD^/T^KO^ at D4 of EB differentiation. Red dots indicate DLK1+ cells. F) Model summarizing molecular regulation of YS and allantois mesodermal lineage specification. Eomes plays a crucial role in differentiation towards the YS mesoderm fate, causing Eomes loss-of-function cells to be redirected towards the allantois lineage. Etv2, required for hematovascular differentiation, is regulated by Eomes during YS formation but not during the formation of allantois endothelium (endo.). The simultaneous KO of Eomes and T, but not their individual KOs, completely ablates the formation of ExEM. Orange bars indicate Eomes expression levels along differentiation trajectories. α; PdgfRa.

Several ExEM populations including the allantois, allantois endothelium and the lateral plate mesoderm form successfully in both Eomes-KO and T-KO chimeric embryos (Fig 3C and 4B). Interestingly, Eomes-KO alters the transcriptomes of allantois and lateral plate mesoderm cells but equivalent cells in T-KO chimeras remain largely unaffected (Fig 4C). The transcriptomes of Eomes-KO and T-KO allantois endothelium, however, are highly similar and not disrupted (Fig 4C). Therefore, disrupting T alone does not impact the formation or the transcriptional properties of YS or allantois cell types, rather neuromesodermal progenitors (NMPs) are the major cell type transcriptionally dysregulated in T-KO chimeras (Fig 4B,C). By contrast, Eomes is required for formation of YS cell types and while Eomes-KO allantois cells have aberrant gene expression profiles, the formation and molecular properties of Eomes-KO allantois endothelial cells appear normal.

Disrupting both Eomes and T simultaneously completely blocks intra-embryonic mesoderm formation in ESC models (Schüle et al., 2023; Tosic et al., 2019), however, the impact on specification of ExEM lineages has not been formally explored. In line with our transcriptional analyses of single KO chimeras, T-KO tdTomato+ and Eomes-KO tdTomato+ ESC contributed to the allantois region in E8.25 chimeric embryos, including Pecam1+ allantoic vasculature (Fig 3D and Fig 4D). In striking contrast, Eomes/T dKO tdTomato+ ESC were absent from both the allantois and YS, indicating a complete block in the ability of Eomes/T dKO ESC to form cell types of these extraembryonic tissues (Figure 4D). Furthermore, depleting Eomes protein in Runx1^Venus^ Eomes^mCherry-degron^ T-KO EBs blocked the production of KDR+ and/or PDGFRA+ mesodermal cell populations (Fig 4E). Together, these findings demonstrate that while individual knockouts of Eomes or T do not disrupt the generation of allantois cells, simultaneous disruption of both T-box factors does.

## Discussion

The specification and diversification of ExEM lineages are crucial for growth and development of the mammalian embryo within the uterine environment. Here, we show a robust EB model mimics temporal induction of ExEM sub-types that arise during gastrulation, generating cell populations that adopt distinct YS or allantoic ExEM fates. Using this system we identify TFs, signaling pathways, and chromatin accessibility changes associated with the emergence of two major ExEM trajectories. Additionally, we identify a mesenchyme population at D4 which potentially represent precursors of two additional ExEM lineages, the amnion and chorion. Our model therefore highlights unexpected mesodermal and endothelial diversity during EB differentiation, thus enabling a more precise interpretation of stem cell differentiation pathways, which will be crucial for potential downstream therapeutic applications.

Using a novel degron-based approach in our EB model, alongside validation experiments in chimeric embryos, we uncover a seminal role for Eomes in directing specification of YS but not allantoic fates. Interestingly, Eomes depletion induces premature activation of TFs and signaling pathways associated with later stages of allantois formation, suggesting Eomes likely represses allantoic ExEM specification in the early posterior PS when YS formation predominates. Notably, our work identifies Eomes as a key regulator of Etv2, aligning with a recent study that demonstrated Eomes overexpression drives upregulation of Etv2 in a similar EB model system (Zhao and Choi, 2017). Previous studies have established Etv2 as an upstream component of the transcriptional hierarchy regulating induction of hematovascular fate in mesodermal precursors in the embryo proper, YS and allantois (Koyano-Nakagawa and Garry, 2017; Koyano-Nakagawa et al., 2012; Liu et al., 2015; Rasmussen et al., 2011). Here, we show that Eomes regulates Etv2 expression at the onset of YS ExEM specification prior to KDR expression, but not during allantois endothelial formation (Fig 4F). Therefore, our research reveals distinct regulatory mechanisms drive Etv2 expression in diverse ExEM lineages and places Eomes at the top of a transcriptional hierarchy specifically regulating YS hematovascular formation.

The T-box family TFs Eomes and T exhibit distinctive expression patterns during murine gastrulation. Eomes is transiently expressed in ExEM precursors that traverse the posterior primitive streak starting at E6.5, while T appears slightly later throughout the primitive streak and nascent mesoderm (Harland et al., 2021). Unlike Eomes, which is crucial for cardiac (Costello et al., 2011) and yolk sac mesoderm specification, T directs mesodermal formation from NMPs, facilitating posterior axis extension (Guibentif et al., 2021; Rashbass et al., 1991; Wilson et al., 1993). Our chimera studies show that neither TF individually have roles in the initial formation of the allantois mesoderm. Previous studies, however, have shown that disrupting both TFs completely blocks intraembryonic mesoderm formation pointing to partial redundancy (Schüle et al., 2023; Tosic et al., 2019). Importantly, our study reveals similar redundancy applying to the extraembryonic compartment, since elimination of both Eomes and T completely ablates ExEM formation both *in vitro* and *in vivo*. Transcriptional disruptions observed in Eomes-KO allantois cells suggest that T can partially compensate for Eomes and drive allantois mesoderm specification. Future research is needed to clarify the partially redundant roles of T-box TFs within ExEM lineages. Our EB model is ideally suited to explore these questions and others, focused on unraveling the genetic networks that specify and pattern extraembryonic tissues—an overlooked yet critical aspect of embryogenesis.

---

**Supplementary Figure 1:**
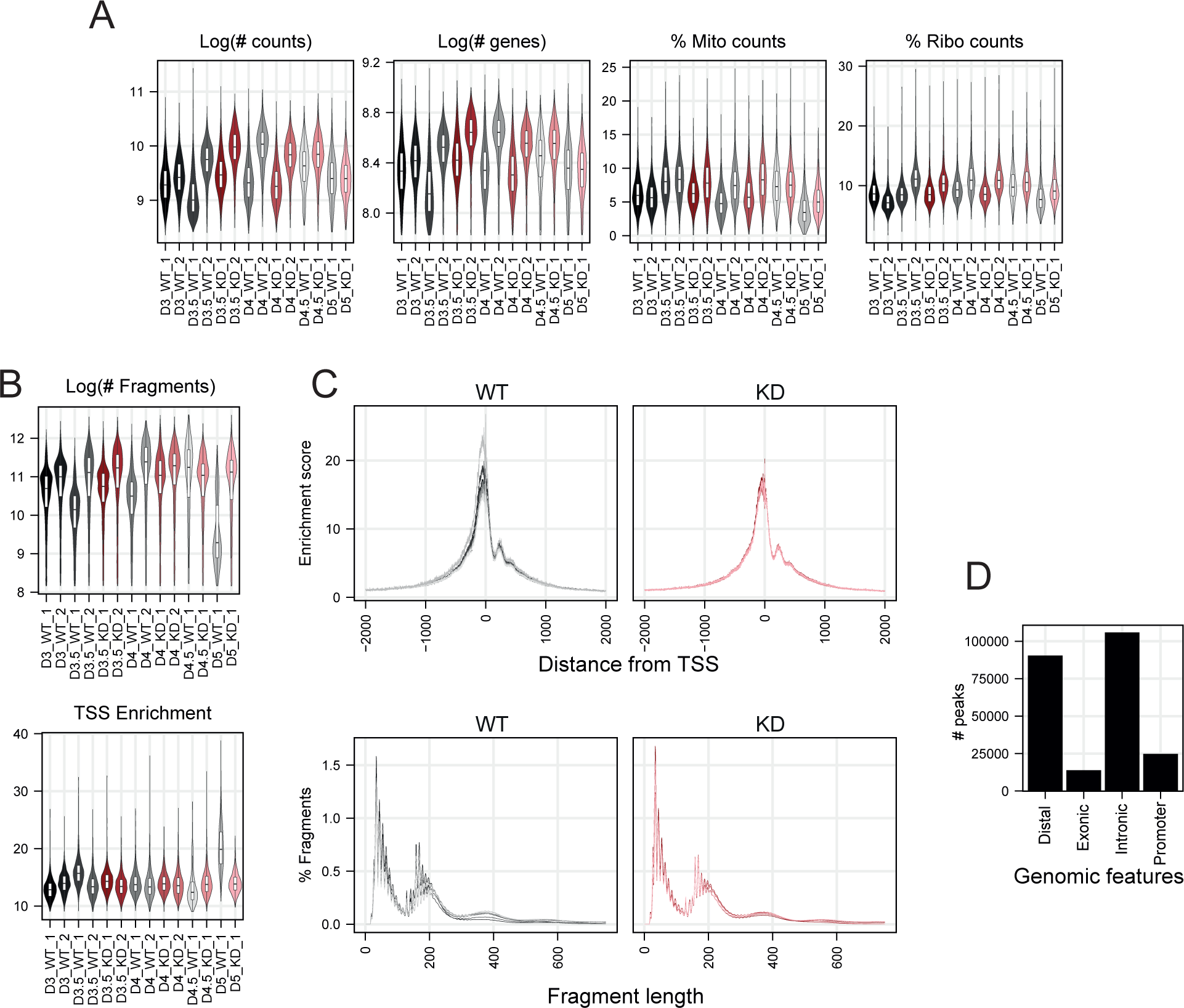
Multiome quality control. A) Violin plots displaying log(number of counts), log(number of genes), percentage of mitochondrial and ribosomal counts per sample. WT and Eomes-KD samples are colored in shades of black and red, respectively. B) Violin plots displaying log(number of fragments) and Transcription Start Sites (TSS) enrichment score, samples colored as in A. C) Genome wide normalized enrichment score around TSS (top) and Fragment length distribution per sample (bottom), samples colored as in A. D) Bar graph displaying number of chromatin peaks per genomic feature.

**Supplementary Figure 2:**
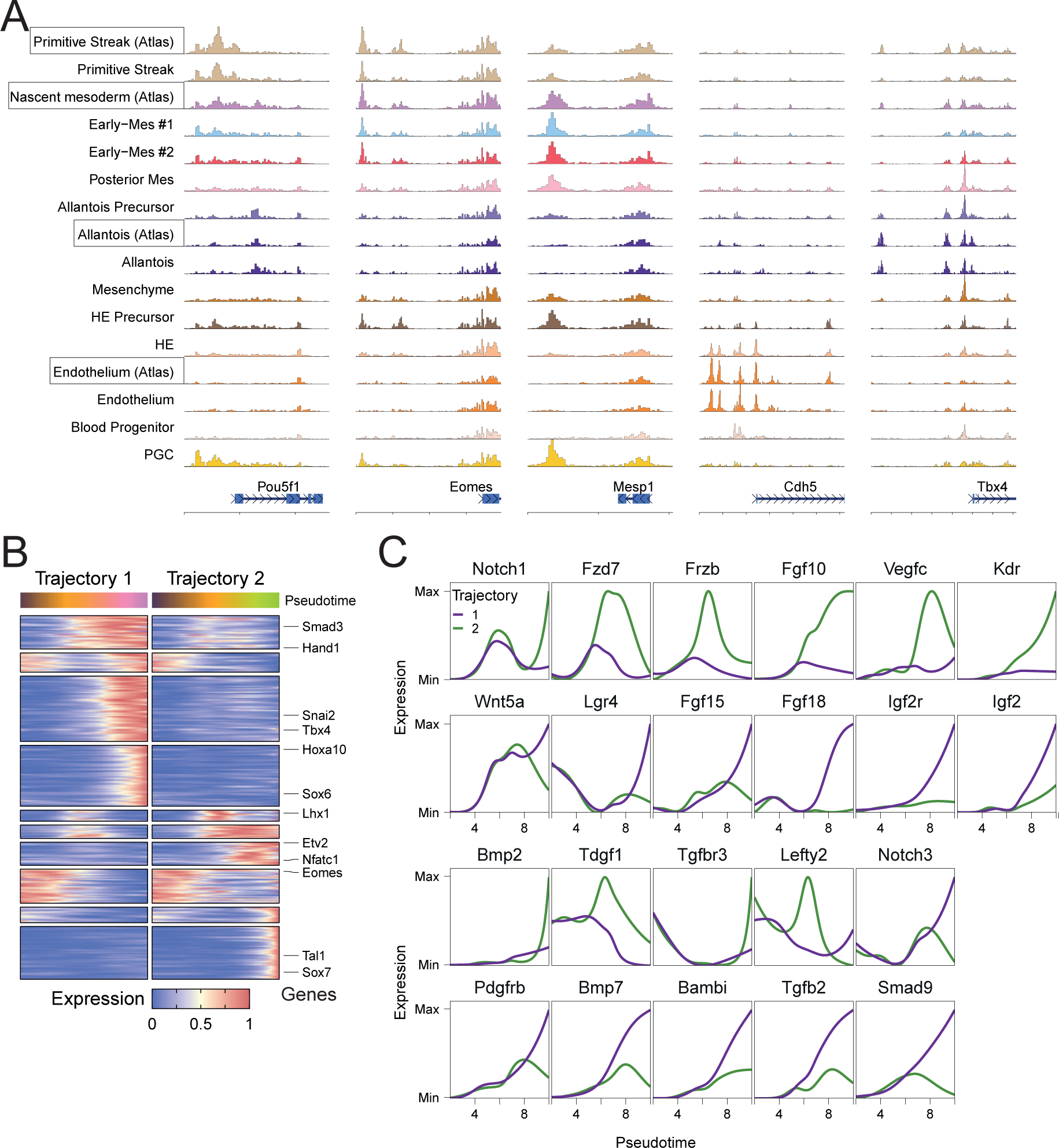
EB differentiation additional information. A) Chromatin accessibility around marker genes for PS (Pou5f1), Early Mesoderm (Eomes/Mesp1), Endothelium (Cdh5), and Allantois (Tbx4). EB cell types are compared to a subset of cell types from a mouse gastrulation multiomic dataset (Atlas) (Argelaguet et al., 2022). B) Heatmap showing normalized gene expression along pseudotime for both trajectories, clustered by expression patterns. Differential testing was performed on pseudotime range indicated in Fig 1F. C) Gene expression (solid lines) for allantois (purple – trajectory 1) and YS (green – trajectory 2) trajectories plotted along pseudotime.

**Supplementary Figure 3:**
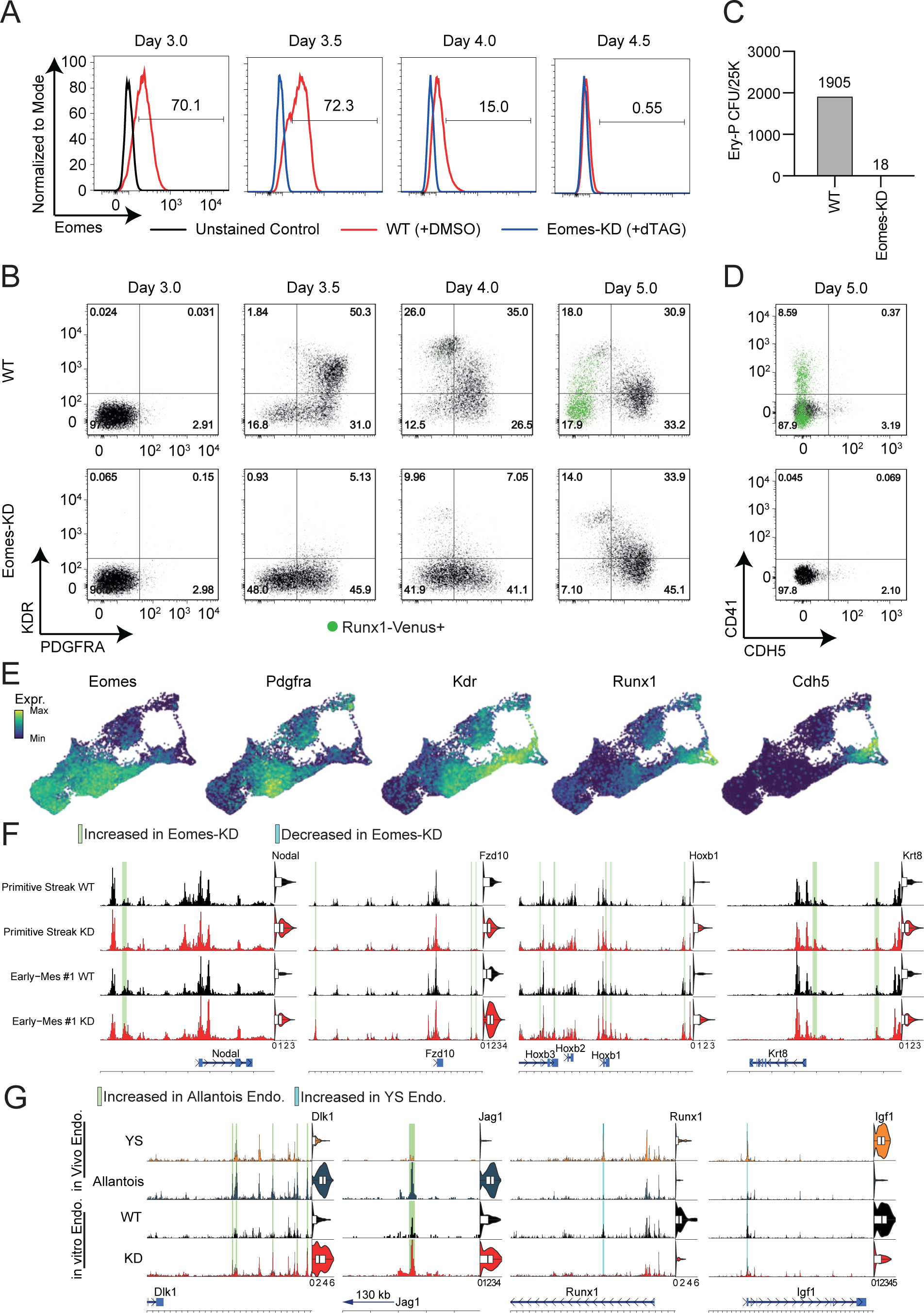
Eomes-KD EB differentiation additional information. A) Intracellular flow cytometric analyses of Eomes protein expression at different timepoints of EB differentiation. The black line at day 3 indicates the signal for the unstained control. EB cultures were treated with DMSO (red lines) and dTAG13 (blue lines) from day 3-4. B) Flow cytometric analyses of KDR and PDGFRA expression at different timepoints of WT (top) or Eomes-KD (bottom) EB cultures. Green dots indicate Runx1-Venus+ cells. C) Primitive erythrocyte colony forming units counted after 5 days of growth in Methocult M3434. Cells were plated at day 5 from single cell suspensions of either WT or Eomes-KD EBs. Ery-P; Primitive Erythrocytes, CFU; Colony Forming Units, 25K; 25,000 cells. D) Flow cytometric analyses of CD41 and CDH5 expression at day 5 of WT (top) or Eomes-KD (bottom) EB culture when DMSO or dTAG13 weas added to cultures from days 3-4. Green dots indicate Runx1-Venus+ cells. E) UMAPs of WT multiome cultures colored by expression levels of indicated genes. F) Genomic region displaying the accessibility and gene expression of a subset of differential genes, with green highlights indicating peaks increased in accessibility in Eomes KD compared to WT in the PS and/or Early-Mes #1. G) Genomic regions displaying the accessibility and expression profile of in vitro Eomes-KD and WT endothelium alongside the *in vivo* allantois and YS endothelium from Argelaguet et al., 2022. Green and turquoise highlights indicate increased and decreased accessibility in Allantois versus YS endothelium, respectively.

**Supplementary Figure 4:**
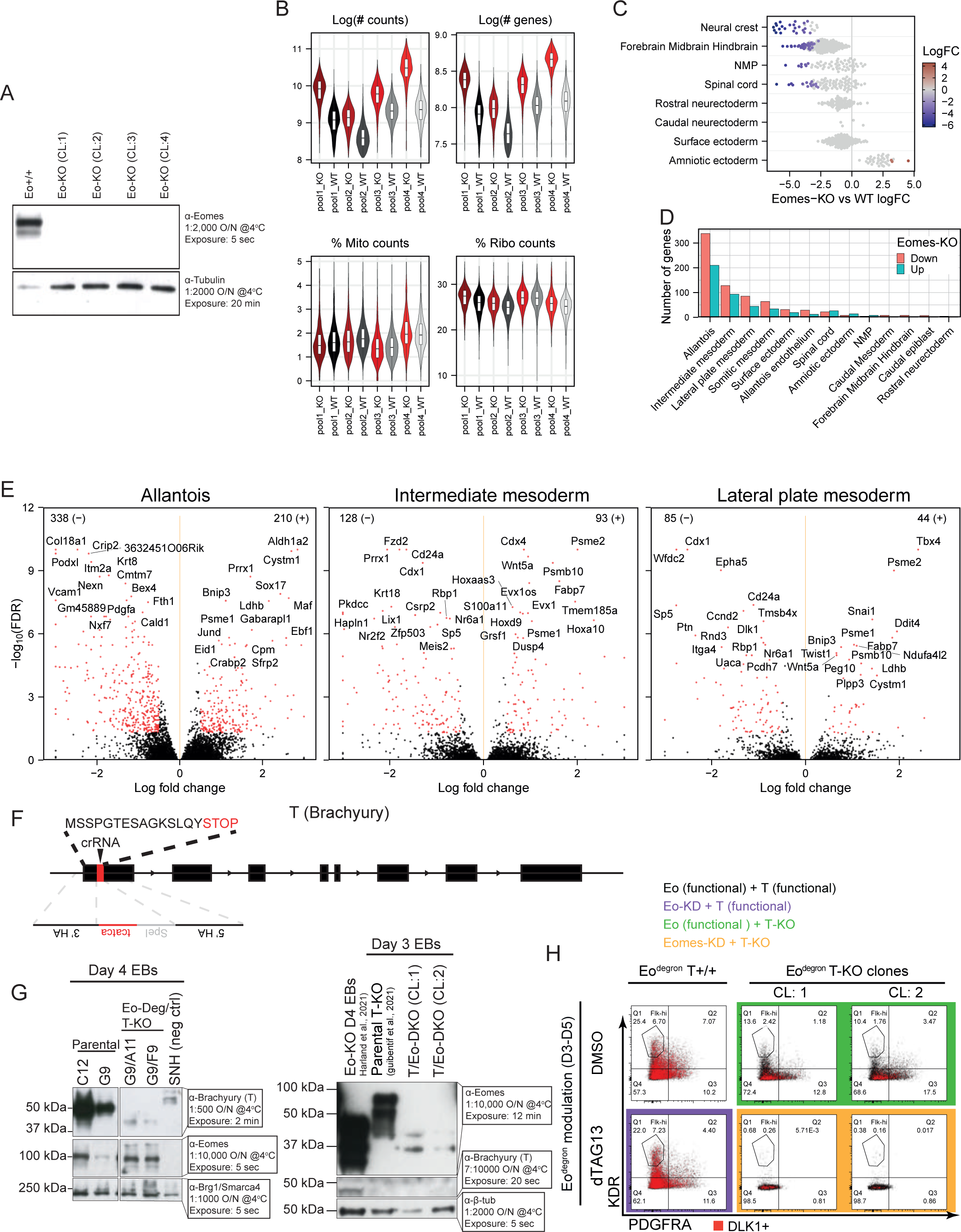
Eomes and T Chimera-seq additional information. A) Western blot for Eomes and α-tubulin expression in tdTomato+ Eomes (Eo) +/+ parental and four KO clones collected at D4 of EB differentiation, confirming Eomes-KO for all four clones used in chimera-seq experiments displayed in Fig 3. B) Violin plots displaying log(number of counts), log(number of genes), percentage of mitochondrial and ribosomal counts per sample of Eomes-KO chimera-seq, with shades of black and red indicating WT host Eomes-KO samples, respectively. C) Differential abundance for Eomes-KO as in Fig 3D, for ectodermal lineages. Negative logFC values indicate depletion of KO cells, while positive logFC values indicate enrichment, neighborhoods without statistical significance are colored grey. D) Number of detected differentially expressed genes that are up (blue) and down (red) regulated in Eomes-KO compared to WT for each cell type. E) Volcano plots for the top three cell types with the most differential genes (allantois, intermediate mesoderm, and lateral plate mesoderm) in Eomes-KO chimera-seq. F) KO strategy for T-KO used for generation of the Eo-Deg/T-KO ESC lines. G) Western blot for T, Eomes, and Smarca4 or α-β-tubulin (loading controls) protein expression collected from D3 and D4 EBs (as indicated) to confirm correct generation of T-KO in Eo-Deg/T-KO (left) and Eomes-KO in Eo/T-dKO lines (right). H) Flow cytometric analyses of PDGFRA and KDR expression for Eomes^WT^/T^WT^, Eomes^KD^/T^WT^, Eomes^WT^/T^KO^, Eomes^KD^/T^KO^ at D5 of EB differentiation. Red dots indicate DLK1 + cells. dTAG or DMSO was added from D3 onwards.

## Methods

### ESC genetic modification using CRISPR-Cas9

#### TdTomato+ Eo-KO and TdTomato+ Eo/T dKO for Chimera-seq

For the generation of TdTomato+ Eo-KO and TdTomato+ Eo/T dKO, two custom crRNAs were used to generate an Eomes loss-of-function allele (Arnold et al., 2008) as described previously (Harland et al., 2021) using TdTomato+ (Pijuan-Sala et al., 2019) and tdTomato+ T-null ESCs lines (Guibentif et al., 2021), respectively. Briefly, ssODNs were designed that contained a 5′ homology arm upstream of the Eomes intron 1 DNA double-strand break site, followed by insertion of a new EcoRV, Sph1 or Spe1 restriction site and a 3′ homology arm located downstream of the Eomes intron 5 DNA break site (Supplementary Table 6). 10 μl of 2 × 10^4^ ESCs in Buffer R were electroporated with 1 μl of a 1:1 mix of the RNPs and 2 μl of the ssODN. Low-density plating was performed after 72 h, and after 7–10 d clones were picked and screened using a three-primer PCR strategy that simultaneously amplified the WT allele and the null allele (Suppl. Table 1). Genotypes of clones were verified using PCR, followed by restriction enzyme digests and Western Blotting.

#### Eo-deg T-KO for EB Differentiations

Feeder-free E14 Runx-Venus Eomes^deg/deg^ ESCs (Bisia et al., 2023) were targeted with a Cas9-mediated strategy similar to that described above, using ssODN template and crRNA guides designed to generate a T-null modification (Suppl. Table 1). Genotypes of clones were verified using PCR, followed by restriction enzyme digests and Western Blotting.

### Western blot

EBs were lysed in RIPA (50 mM Tris pH8.0, 150 mM NaCl, 1% Igepal, 0.5% Na deoxycholate, 0.1% SDS). Protein quantification by Bradford was carried out using DC^TM^ Protein Assay Kit I (BioRad) on a Jenway Genova DNA Life Science Analyzer. Samples were denatured at 98^ο^C for 10 min in Laemmli buffer (BioRad) with 10% β-mercaptoethanol, run on a Mini-Protean® PAGE gel (BioRad) at 90V and transferred onto PVDF membrane for 75 min at 90V. The membrane was rinsed with dH2O, washed in 0.1% TBST (Tris-buffered saline-Tween20) for 10 min, blocked with EveryBlot blocking buffer (BioRad) for 10 min at room temperature incubated with primary antibody on a shaking platform overnight at 4^ο^C, washed with TBST, incubated in secondary antibody, and washed with TBST. ECL prime (Sigma-Aldrich) was added per manufacturer’s instructions and exposed to X-ray film. When required, the membrane was stripped with stripping buffer (2.9g glycine, 20ml SDS in 2l H2O, pH2.2), blocked, washed, and exposed to antibody as above. Antibodies used are listed in Suppl. Table 1.

### ESC maintenance

All TdTomato+ mouse ESC lines were cultured under 2i+LIF conditions (Ying et al., 2008) prior to blastocyst injection for chimera-seq experiments. Runx1^Venus^ Eomes^mCherry-degron^ ESC lines were maintained in feeder free culture conditions in serum + LIF as previously described (Costello et al., 2011).

### In vitro hematovascular EB differentiation protocol

48-72 hrs prior to induction of hematovascular differentiation ESC were washed with PBS and cultured in serum free ESC media containing 50% Neurobasal Media (Gibco, Cat #21103049), 50% DMEM/F12 (Gibco, Cat#11320033), supplemented with 0.5X of both N2 (Gibco, Cat #17502048) and B27 (Gibco, Cat #17504044), 1%Pen/Strep, 1% glutamine, 0.05% BSA (Gibco, Cat #15260037), 1 uM PD0325091, 3 uM CHIR99021 and 1000 U/ml LIF. At D0 cells were dissociated using Trypsin-LE (Gibco) and seeded at a density of 1×10^5 cells ml in serum-free differentiation (SF-D) media (Nostro et al., 2008) and cultured on an orbital shaker at 70 rpm for ∼18 hrs in the absence of growth factors to form EBs. At D2, EBs were split 1:3 in SF-D media containing recombinant human (rh) VEGF (5 ng ml-1; R&D Systems), rhBMP4 (10 ng ml-1; R&D Systems) and Activin A (5 ng ml-1; R&D Systems) for 48 hr. At D3, EBs were treated with either DMSO for WT controls or dTAG-13 for Eomes-KD (100 nM, Biotechne Tocris Cat #6605). At D4, EBs were split 1:2 in SF-D media containing rhVEGF (5 ng ml-1; R&D Systems), rhBMP4 (10 ng ml-1; R&D Systems) and Activin A (5 ng ml-1; R&D Systems) for 24 hr.

### Flow cytometry

#### Live flow cytometry

For live flow cytometry, EBs were washed with PBS, dissociated with TrypLE and neutralised with FACS buffer (PBS with 1% Pen/Strep, 2% FCS (fetal calf serum)). Cells were resuspended in FACS buffer containing fluorophore-conjugated antibodies and stained on ice for 30 min. Cells were washed and resuspended in FACS buffer with 1:5,000 DAPI (BD Biosciences) for 15 min on ice, washed and resuspended again in FACS buffer. Antibodies used are listed in Suppl. Table 1.

#### Intracellular flow cytometry

For intracellular flow cytometry, EBs were washed with PBS, dissociated with TrypLE and neutralised with FACS buffer (PBS with 1% Pen/Strep, 2% FCS (fetal calf serum)). Next, cells were stained with an eBioscience™ Fixable Viability Dye eFluor™ 450 (Ref: 65-0863-14) in a 96-well plate for approximately 30 minutes, washed with PBS and subsquently fixed and permeabilized using the eBioscience™ Foxp3/ Transcription Factor Staining Buffer Set (Ref: 00-5523-00) following manufacturer instructions. Cells were stained with an anti-Eomes AF88 antibody (Ref: 53-4875-82) at a concentration of 1:400 for 30 minutes at room temperature, washed and resuspended again in FACS buffer for flow cytometric analyses.

### Collecting EB samples for 10x multiomics and flow cytometry

In two separate experiments (EB differentiations performed weeks apart) live and intracellular flow cytometry (SFig3), as well as 10x multiomics (Fig1 and 2), were performed on D3, D3.5, D4, D4.5 and D5 EB samples. The start time of the differentiation setup was staggered, allowing for the collection and dissociation of EBs at intervals of days 3, 3.5, 4, 4.5, and 5 from both DMSO and dTAG treated conditions on the same collection day. Each sample was split such that half of the cells were taken for 10x multiomics and the other half for live and intracellular flow cytometry.

### Nuclear extraction and 10x genomics multiome library preparation

Cell suspensions in 2mL eppendorf tubes were centrifuged at 300g for 5 minutes, resuspended in 1 mL of PBS supplemented with 0.04% BSA then filtered using a 40uM flowmi cell strainer. After centrifuging again, the supernatant was removed and the cell pellet resuspended in 100ul of ice cold nuclear extraction (NE) buffer (10mM Tris pH 7.5, 10mM NaCl, 3mM MgCl2, 1% BSA, 0.1% Tween, 1mM DTT, 1U/ul RNaseIn (Promega), 0.1% NP40, 0.01% Digitonin) and incubated on ice for 4 minutes. 1mL of wash buffer (identical to NE buffer but lacking NP40 and digitonin) was added and nuclei were centrifuged at 500g for 5 minutes at 4°C. Nuclei washed twice by resuspension in 1mL of wash buffer and centifugation at 500g for 5 minutes at 4°C. Nuclei were resuspended in 50ul of diluted nuclei buffer (10x Genomics) and 1ul was used to assess quality using a microscope and count nuclei using a Countess II instrument. >99% of nuclei stained positive for trypan blue and the nuclei were found to have the expected morphology. Nuclei were diluted such that a maximum of 16,000 were taken forward for 10x Multiome library preparation. Libraries were prepared using the 10x Genomics Chromium and sequenced on a Novaseq 6000 instrument (Illumina) using the recommended read-lengths.

### RNA+ATAC Multiome analysis

#### Mapping sequencing data

10x genomic multiome raw base call BCL were first demultiplexed using cellranger-arc mkfastq (Cell Ranger ARC version 2.0.1) in order to get separate FASTQ files for the GEX (RNA) and ATAC libraries. Next, FASTQ files were mapped to the mm10 reference genome supplied by 10x Genomics (arc-mm10-2020-A-2.0.0) using cellranger-arc count. All downstream processing and analysis were performed in R (v4.2.2). Analysis started with the pre-processing of the RNA data, followed by pre-processing the ATAC and analysis of the combined modalities.

### scRNA-seq pre-processing

From the scRNA-seq, high quality cells were retained using the following thresholds: log10(number of reads) > 3 & < 5, number of genes > 2.5e3 & < 12e3, percentage mitochondrial RNA < 25%, percentage ribosomal RNA < 30%. Count normalization was performed by first calculating size factors using the computeSumFactors function from scran ((Lun et al., 2016) Version 1.26.0), where cells were pre-clustered using scran’s quickCluster with method=igraph, minimum and maximum sizes of 100 and 3000 cells per cluster, respectively, followed by logNormCounts from scuttle ((McCarthy et al., 2017) Version 1.8.0). Doublet calling was performed with the scds ((Bais and Kostka, 2020) Version 1.14.0) package using the hybrid approach and doublet score > 1.25 as threshold. A total of 35,125 cells passed RNA QC.

### scRNA label transfer from the scRNA-seq atlas of gastrulation and early organogenesis

Label transfer for cell-type assignment was conducted by mapping to the E6.5 – E8.5 stages of the gastrulation atlas, with additional cell type annotation from an updated extended gastrulation atlas (Imaz-Rosshandler et al., 2024; Pijuan-Sala et al., 2019). First, the reference atlas was subset to a maximum of 15,000 cells per embryonic stage, including all cells for the ‘mixed-gastrulation’ stage. Mapping was performed for each sample separately using the batchelor package (Version 1.14.0)(Haghverdi et al., 2018), with each sample referred to as the ‘query’. For mapping to the reference atlas, non-informative and noisy genes were excluded, including those with names starting or ending with Rik, Mt, Rps, Rpl, or Gm, as well as haemoglobin genes, imprinted genes Grb10 and Nnat. Subsequently, the top 2,500 genes with the highest variability in expression across atlas celltypes were selected for integration. Joint normalization was performed using multiBatchNorm with batch being either ‘query’ or ‘atlas’, followed by multiBatchPCA with the same batch variable and d = 50. Batch correction was executed in multiple rounds using ReducedMNN. Initially, the atlas was corrected within each embryonic stage, ordering samples from largest to smallest. This was followed by correcting the atlas between embryonic stages, ordered from latest to earliest. Lastly, batch correction was applied between the reference atlas and the query. Nearest neighbors (NN) of each query cell in the reference atlas were identified using queryKNN from the BiocNeighbors package (Version 1.16.0) with k=25. Cell-type label transfer was then performed by determining the mode of the cell types of the NN, with ties resolved by assigning the cell type of the reference atlas cell closest to the query.

### scRNA dimensionality reduction

Dimensionality reduction was performed on lognormalized counts by first calculating the top variable genes using Seurat’s (Version 4.3.0) FindVariableFeatures with nfeatures = 1500 after excluding the same non-informative and noisy genes as described above. Next, the number of reads and genes, as well as the percentage of mitochondrial and ribosomal reads were regressed out for the lognormalized counts. PCA was then performed using scater’s ((McCarthy et al., 2017), Version 1.26.0) runPCA with ncomponents = 15.

### scATAC pre-processing

Full scATAC-seq analysis was performed using the ArchR package (Vversion 1.0.1)(Granja et al., 2021). Arrow files were created from CellRangers fragment files using createArrowFiles using initial relaxed quality threshold for the transcription start site enrichment (min/max TSS) and number of frags (min/max Frags): minTSS = 2.5, minFrags = 1e3, and maxFrags = 1e6, while excluding chromosomes ‘chrM’ and ‘chrY’, followed by creation of the ArchR object using the ArchRProject function for the arrow files from all samples. After inspecting quality, cells were retained using the following, more stringent, thresholds: minTSS = 9, maxTSS = 35, and minFrags = 3.5e3, as well as a minimum proportion of reads in blacklisted genomic regions of 0.05. Cells detected as doublets in the RNA were excluded from downstream analysis, while cells that were only detected in the ATAC but not in the RNA, while passing ATAC QC thresholds, were kept. A total of 35,864 cells passed ATAC QC.

### scATAC peak detection and dimensionality reduction

The mouse genome was split into 500 bp bins in order to create a count matrix with accessibility per bin for each cell using ArchR’s addTileMatrix. Next, Latent Semantic Indexing (LSI) dimensionality reduction was performed on the TileMatrix using addIterativeLSI with iterations = 4, varFeatures = 2e4, dimsToUse = 1:30, and clusterParams’ resolution = c(0.4, 1, 2) and maxClusters = NULL. The LSI was used for clustering data using addClusters with resolution = 0.5. Next, insertion coverage files were creates for pseudo-bulked replicates of each TileMatrix LSI cluster and Eomes condition (KD or WT) combination using addGroupCoverages with groupBy = the cluster & condition combination, minCells = 50, maxCells = 5000. These insertion coverage files were used for Macs2 peak calling using the function addReproduciblePeakSet with groupBy = the cluster & condition combination, cutOff = 1e-3, and extendSummits = 300, for peaks with a width of 600 bp. A count matrix for the accessibility of those peaks per cell was created using addFeatureMatrix with the Granges object of the peaks as features, and ceiling = 4. LSI dimensionality reduction was performed on the PeakMatrix in the same way as described above for the TileMatrix, but with varFeatures = 2.5e4, dimsToUse = 1:20.

### *In-silico* ChIP of the *in vivo* multiome atlas of murine embryology

In order to predict TF occupancy at each of the identified peaks from the *in vitro* multiome, we implemented the *in silico* ChIP method on the *in vivo* multiome gastrulation atlas as previously described (Argelaguet et al., 2022). Briefly, for all cells that pass quality thresholds in both the RNA and ATAC modality of the *in vivo* data, we first identified high resolution clusters needed to correlate TF expression with peak accessibility. Transcriptomics counts were lognormalised using Scuttle’s logNormCounts, followed by identification of the top 5000 highly variable genes, ordered by FDR, using Scran’s modelGeneVar function with samples as block. Next, PCA was performed using runPCA with ncomponents = 40 on all HVGs. High resolution clustering was then performed using seurat’s functions FindNeighbors across all PC dimensions and FindClusters with a resolution of 20 in order to identify 222 clusters. These clusters were used to pseudobulk the count matrices for the RNA expression and the ATAC accessibility of the *in vitro* peak set using Scuttle’s aggregateAcrossCells. Pseudobulk matrices were normalized by dividing counts by total counts per cluster, multiplying by 1e6 and adding a pseudocount of 1, followed by taking the natural log. Next, motif detection in the *in vitro* peaks was performed using the cisbp dataset of the chromVARmotifs package (Version 0.2.0). To make motif names consistent with gene names, the Tcfap family was renamed to tfap. Motif annotation was performed using motifmatchr’s (Version 1.20.0) matchMotifs function with out = ‘scores’, p.cutoff = 5e-5, and w = 7. Next, for every TF, we used psych’s (Version 2.2.9) corr.test function to calculate the pearson correlation between the normalized counts of its expression with the normalized counts of the accessibility of peaks containing its respective motif. The *in silico* binding score for each TF to a peak was calculated by multiplying this correlation with the minmax normalized product of the TF motif score of the peak and the maximum normalized accessibility of the peak across all clusters. An *in silico* binding score of > 0.2 was set as a threshold for a peak to be predicted as ‘bound’ by a specific TF. The *in silico* ChIP motif annotation is used for any motif analysis throughout the paper.

### scATAC ChromVAR

In order to calculate TF activity on a per-cell basis, we employ ArchR’s ChromVAR implementation using the addDeviationsMatrix function, with the matches being the peak annotation for motifs with a predicted binding score of 0.2. TFs with fewer than 40 peaks predicted to be bound were excluded from downstream analysis.

### scRNA & scATAC integration

In order to integrate the scRNA and scATAC, we used the PCA and LSI calculated from the RNA and ATAC, respectively, scaled each component to achieve comparable ranges for both PCA and LSI, and performed MOFA+ (Version 1.8.0)(Argelaguet et al., 2020) using default settings on the subset of cells that passed QC for both modalities. This resulted in a total of 19 MOFA factors that were used for downstream analysis. UMAP was calculated from MOFA factors using uwot’s (Version 0.1.14) umap function with n_neighbors = 25 and min_dist = 0.15. Clustering was performed on MOFA factors using Seurat’s FindNeighbors with k.param = 35, followed by FindClusters with resolution = 1.5 in order to identify a total of 18 clusters. Using a combination of marker genes, marker motifs, and cell type label transfer from the in vivo atlas, we labelled the 12 cell types used for downstream analysis throughout the paper. Bigwigs for the 12 cell types were generated using ArchR’s getGroupBW with default parameters

### Wild-type analysis

#### Cell type specific markers

For the analysis of the WT EBs marker genes for the 12 clusters were identified using Seurat’s FindAllMarkers with min.pct = 0.1, logfc.threshold = 0.5, and only.pos = TRUE. ChromVAR marker motifs were identified using ArchR’s getMarkerFeatures with useSeqnames="z", with Fig 1D showing -log10(FDR) displayed as the size and chromVAR Z-scores minmax normalized per motif.. Marker peaks were identified using ArchR’s getMarkerFeatures, followed by motif enrichment using peakAnnoEnrichment with cutOff = "FDR <= 0.1 & Log2FC >= 0.5" and plotting data was retrieved using plotEnrichHeatmap with returnMatrix = TRUE.

#### Trajectory inference and differential testing

Trajectory inference was performed using the slingshot package (Version 2.6.0)(Street et al., 2018). Briefly, the data was limited to the celltypes of Primitive Streak, Early Mes #1 and #2, Posterior Mesoderm, Allantois Precursor, HE Precursor, and HE. Slingshot trajectories and pseudotimes were calculated using the slingshot function, with celltypes as clusterLabels, the ‘Primitive Streak’ as start.clus, and the first 10 dimensions of the MOFA factors as reducedDim. Cells were assigned to either the allantois or YS trajectory using the .assignCells function of the tradeSeq package (Version 1.12.0)(Van den Berge et al., 2020), with the slingshot weights as input. Differential testing was restricted to WT cells within pseudotime 2 and 10, and to the top 1e4 highly variable genes. Differential testing along pseudotime was performed by first running Tradeseq’s fitGAM, with the described cells and slingshot pseudotime, the hvgs as genes, nknots = 8, and with the multiome replicates used as the design matrix. Next, differential genes were detected by using patternTest and diffEndTest, both with l2fc = 1, and genes with an FDR < 0.05 were labelled as significant. For the differential chromatin accessibility, fitGAM was run using the same parameters, with the exception that the count matrix was replaced by the ATAC count matrix and the hvgs were the top 42261 most variable peaks. patternTest and diffEndTest were used with l2fc = 0.25 and peaks with an FDR < 0.05 were labelled as significant. Motif enrichment for peak clusters was performed using ArchR’s .computeEnrichment.

### Knock-down analysis

#### Differential abundance testing

Differential abundance testing was performed at a single-cell level after excluding the day 3 WT cells. Briefly, for each timepoint we determined the global expected KD ratio (KD cells/total cells). Next, for each cell, we identified the top 100 nearest neighbours using BiocNeighbors’ findKNN function using the mofa factors. For each cell we then calculate the per cell expected KD ratio by taking the sum of global expected KD ratios normalized for the number of neighbours belonging to each timepoint and dividing by 100. P values were calculated by performing a two-sided binomial test with a confidence level of 0.95, testing if the observed KD ratio is different from the per cell expected KD ratio, followed by FDR multiple testing comparison. Levels of differential abundance were defined as the per cell observed divided by the per cell expected KD ratio, as displayed in Fig 2B.

#### Differential genes and accessibility in early cell types and endothelium

Differential expression and accessibility testing between KD and WT cells was performed using edgeR (Version 3.40.0)(Chen et al., 2016) on pseudobulk replicates. Briefly, for each cell type – genotype condition we generated 5 pseudobulk replicates by randomly sampling 40% of cells for each replicate. Genes were filtered by setting minimum thresholds for expression levels and cellular detection rates and a set of non-informative genes were removed before running edgeR identify DEGs between Eomes KD and WT. Significance was determined as an absolute logFC > 0.5 and FDR < 0.05. Differentially accessible regions were determined in the same way, using the peak count matrix rather than the RNA expression count matrix.

#### Gastrulation multiome atlas endothelium annotation refinement

In order to improve the resolution of annotation for the endothelial subsets of the multiome gastrulation atlas (Argelaguet et al., 2022) the subset of ‘Haematoendothelial_progenitors’ and ‘Endothelium’ were re-analyzed. Dimensionality reduction was performed on the top 3000 highly variable genes, identified using scran’s modelGeneVar, followed by batchelor’s multiBatchPCA with samples as batch and d = 10. Cells were then clustered using Seurat’s FindNeighbors with k = 20 and FindClusters with resolution = 0.3. Clusters were then annotated by comparing the expression of cluster marker genes with previously published genes marking endothelium from the different anatomical regions (Imaz-Rosshandler et al., 2023).

### Eomes chimera generation

tdTomato-ESC clones used for chimera generation included Eomes-KO (This study), T-KO (Guibentif et al., 2021), Eomes/T-dKO cells (this paper) and wild-type controls (Pijuan-Sala et al., 2019). E3.5 blastocysts were derived from wildtype C57BL/6 matings, injected with 4-6 ESC and transferred into 2.5 day pseudopregnant recipient females. Chimeric embryos were harvested at E8.5, dissected and processed for either imaging or scRNA-seq. Embryos for scRNA-seq were dissociated using TrypLE Express dissociation reagent (Thermo Fisher Scientific) incubation for 7-10 minutes at 37°C under agitation followed by sorting of single-cell suspensions into tdTomato+ (KO) and tdTomato- (WT) samples using a BD Influx sorter with DAPI at 1mg/ml (Sigma) and subsequent 10X single-cell 3’ RNA-seq. 4 separate Eomes-KO clones were used and each replicate consists of multiple embryos pooled together from individual clones.

### Chimera-seq analysis

#### Mapping sequencing data and *Pre-processing*

Eomes 10x genomic scRNA-seq FASTQ files were mapped to the mm10 reference genome and counted with the GRCm38.p5 annotation including the Tomato-Td gene using CellRanger count (v6.0.1) with chemistry=SC3Pv3. T chimera data for reanalysis was downloaded using the package MouseGastrulationData (Version 1.12.0) with the function TChimeraData with type = ‘processed’ and samples = c(1:2, 5:10). Eomes and T chimera scRNA-seq analysis was performed with the same parameters unless indicated. Chimera scRNA-seq pre-processing was performed in the same way as described above for the scRNA of the multiome assay, with a few exceptions. The thresholds were adjusted to log10(number of reads) > 3.5 & < 5, number of genes > 1.5e3 & < 1e4, percentage mitochondrial RNA < 5%, percentage ribosomal RNA < 35%. Doublet detection was performed as described, with a threshold of > 0.5 set as a doublet call. For the newly generated Eomes chimera-seq, a total of 22,600 WT and 9768 Eomes-KO cells were retained for downstream analysis. For the re-analysis of the E8.5 T chimera-seq, a total of 13,771 WT and 14,460 T-KO were retained.

#### Mapping and refined cell type annotation

Chimera scRNA-seq mapping to the reference atlas was performed in the same way as described above for the scRNA of the multiome assay in the ‘scRNA label transfer from the scRNA-seq atlas of gastrulation and early organogenesis’ section, with a few exceptions. Query cells were only mapped to atlas cells spanning stages E7.5 to E8.5. Additionally, genes starting or ending with Rik, Mt, Rps, Rpl, or Gm, haemoglobin genes, imprinted genes, sex related genes Xist, Tsix, and y-chromosomal genes, and td-Tomato itself were excluded for mapping and multiBatchPCA was performed on the top 4000 most variable genes in the atlas with d = 40. After batch correction, the label of the 15 NN of the reference were used for label transfer. In order to increase the resolution in annotation for the cell types of interest, relevant labels from the original and extended gastrulation atlas were combined and used for downstream analysis (Imaz-Rosshandler et al., 2024; Pijuan-Sala et al., 2019).

#### Milo differential abundance testing

Differential abundance testing was performed for each embryonic stage separately using the MiloR package (Version 1.6.0)(Dann et al., 2021). Cells with an annotation of "Visceral_endoderm", "ExE_endoderm", "ExE_ectoderm", or "Parietal_endoderm" were removed from the analysis. Dimensionality reduction was performed by first selecting the top 4000 highly variable genes of the WT samples, identified by Seurat’s FindVariableFeatures, excluding genes starting or ending with Rik, Mt, Rps, Rpl, or Gm, haemoglobin genes, imprinted genes, sex related genes Xist, Tsix, and y-chromosomal genes, and td-Tomato itself. Next, the number of reads and genes, as well as the percentage of mitochondrial and ribosomal reads were regressed out for the lognormalized counts. Principle component analysis was performed using multiBatchPCA with the genotype set as batch variable and d = 40, followed by batch correction using reducedMNN. Next, Milo was performed on batch corrected PCs by first running buildGraph with k = 20 for Eomes and k = 40 for T chimeras, followed by makeNhoods with the same values for k and prop=0.1. Cells per neighbourhood were counted using countCells and differential abundance testing was performed using testNhoods with design = ∼ embryo pool + genotype, where each embryo pool contains one sample for both the KO and the WT condition that were collected as chimaeric embryos and separated by the presence or absence of tdTomato expression by fluorescence-activated cell sorting.

#### Differential gene expression testing

Differential gene expression testing was performed by first aggregating cells by their sample, genotype, and cell type annotation. Then, using scran’s pseudoBulkSpecific with design = ∼ embryo pool + genotype, we identified genes that were differentially expressed between the KO and WT cells in a cell type specific way, with p-values being calculated by comparing gene specific logFC to the average logFC across all cell types. Significance was determined as an absolute logFC > 0.5 and FDR < 0.05. For comparison between T and Eomes DEGs, genes that were excluded from one of the cell types in one of the KOs were set as a logFC of 0.

### Chimera imaging

Embryos were washed in PBS and fixed in 4% paraformaldehyde overnight at 4oC. After three washes in PBS containing 0.1% Triton X-100 (PBS-T), samples were permeabilised in PBS containing 0.5% Triton X-100 at RT for 20mins, followed by further washes in PBS-T. Embryos were blocked in 5% donkey serum, with 0.2% BSA in PBS-T at RT for 2 hours then incubated overnight with primary antibodies in blocking buffer at 4oC. After multiple washes in PBS-T, embryos were incubated with fluorophore-conjugated secondary antibodies in blocking buffer for 2 hour at RT, followed by further washes in PBS-T, including a PBS-T/DAPI wash. Embryos were mounted in Vectashield with DAPI on coverslip dishes and imaged on an Olympus Fluoview FV1000 microscope and image data were processed using ImageJ software. Antibodies are listed in Suppl. Table 1.

## Acknowledgements

We thank William Mansfield at the CRUK Pre-clinical Genome Editing facility for blastocyst injections, the Flow Cytometry Core Facility at CIMR for cell sorting, CRUK-CI genomics core for preparing the single-cell libraries, the Wellcome Sanger Institute DNA Pipelines Operations for sequencing, Darran Clements from the CSCI Imaging Facility for microscopy support, Rebecca Hannah for data support, and the Cambridge Research Computing Services for supplying computer resources. We also thank the Dunn School Don Mason Flow Cytometry Facility, the Dunn School Bioimaging Facility, Genome Engineering Oxford, and the Pathology Support Building Facility. Finally, we would like to thank all members of the Robertson and Gottgens labs for their advice and support, including Philipp Kurbel and Atreyi Biswas for technical assistence.

## Author contributions

Conceptualization and research design: L.T.G.H, B.T., E.J.R., B.G.; EB differentiation experiments: L.T.G.H, A.B.; Bioinformatic analyses: B.T.; ESC cell-line generation: L.T.G.H, A.B., E.J.R.; 10X multiomics sample processing: T.L., S.C.; Chimera injections and dissections: M-L.N.T, E.J.R., I.C.; Imaging: I.C.; Project support: N.W., E.B; Data curation: B.T., R.A; Writing: L.T.G.H, B.T, B.G, E.J.R wrote the manuscript with feedback from co-authors. Supervision: E.J.R, B.G.; Project administration: L.T.G.H, E.J.R, B.G., W.R.; Funding acquisition: L.T.G.H, B.T., B.G., E.J.R., A.M.B.

## Data and code availability

Data from this study have been deposited at the Gene Expression Omnibus (GSE274166, GSE274167). Previously published T-KO chimera scRNA-seq was reanalysed and can be found on Arrayexpress under code E-MTAB-8811 or accessed via the R package ‘MouseGastrulationData’ via the ‘TChimeraData’ function. Reference atlas data used throughout the paper can be accessed at https://marionilab.github.io/ExtendedMouseAtlas/ for scRNA-seq (Imaz-Rosshandler et al., 2024; Pijuan-Sala et al., 2019) and GSE205117 for the multiome gastrulation atlas (Argelaguet et al., 2022). All code used to process and analysed the data is available at https://github.com/BartTheeuwes/Eomes_ExEM.

## Declaration of generative AI and AI-assisted technologies in the writing process

During the preparation of this work the author(s) used ChatGPT in order to improve clarity of writing. After using this tool/service, the author(s) reviewed and edited the content as needed and take(s) full responsibility for the content of the publication.

## References

Argelaguet, R., Arnol, D., Bredikhin, D., Deloro, Y., Velten, B., Marioni, J. C. and Stegle, O. (2020). MOFA+: a probabilistic framework for comprehensive integration of structured single-cell data. Genome Biol. 837104.

Argelaguet, R., Lohoff, T., Li, J. G., Nakhuda, A., Drage, D., Krueger, F., Velten, L., Clark #, S. J. and Reik #, W. (2022). Decoding gene regulation in the mouse embryo using single-cell multi-omics. bioRxiv 2022.06.15.496239.

Arnold, S. J., Hofmann, U. K., Bikoff, E. K. and Robertson, E. J. (2008). Pivotal roles for eomesodermin during axis formation, epithelium-to-mesenchyme transition and endoderm specification in the mouse. Development 135, 501–511.

Arora, R. and Papaioannou, V. E. (2012). The murine allantois: A model system for the study of blood vessel formation. Blood 120, 2562–2572.

Bais, A. S. and Kostka, D. (2020). Scds: Computational annotation of doublets in single-cell RNA sequencing data. bioRxiv 36, 1150–1158.

Bisia, A. M., Costello, I., Xypolita, M.-E., Harland, L. T. G., Kurbel, P. J., Bikoff, E. K. and Robertson, E. J. (2023). A degron-based approach to manipulate Eomes functions in the context of the developing mouse embryo. Proc. Natl. Acad. Sci. 120, 1–10.

Chen, Y., Lun, A. T. L. and Smyth, G. K. (2016). From reads to genes to pathways: Differential expression analysis of RNA-Seq experiments using Rsubread and the edgeR quasi-likelihood pipeline. F1000Research 5, 1–51.

Ciruna, B. G. and Rossant, J. (1999). Expression of the T-box gene Eomesodermin during early mouse development. Mech. Dev. 81, 199–203.

Costello, I., Pimeisl, I. M., Dräger, S., Bikoff, E. K., Robertson, E. J. and Arnold, S. J. (2011). The T-box transcription factor Eomesodermin acts upstream of Mesp1 to specify cardiac mesoderm during mouse gastrulation. Nat. Cell Biol. 13, 1084–1092.

Dann, E., Henderson, N. C., Teichmann, S. A., Morgan, M. D. and Marioni, J. C. (2021). Differential abundance testing on single-cell data using k-nearest neighbor graphs. Nat. Biotechnol. 2021 1–9.

Ditadi, A., Sturgeon, C. M. and Keller, G. (2017). A view of human haematopoietic development from the Petri dish. Nat. Rev. Mol. Cell Biol. 18, 56–67.

Downs, K. M. (2022). The mouse allantois: new insights at the embryonic-extraembryonic interface. Philos. Trans. R. Soc. B Biol. Sci. 377,.

Dzierzak, E. and Bigas, A. (2018). Blood Development: Hematopoietic Stem Cell Dependence and Independence. Cell Stem Cell 22, 639–651.

Granja, J. M., Corces, M. R., Pierce, S. E., Bagdatli, S. T., Choudhry, H., Chang, H. Y. and Greenleaf, W. J. (2021). ArchR is a scalable software package for integrative single-cell chromatin accessibility analysis. Nat. Genet. 53, 403–411.

Guibentif, C., Griffiths, J. A., Imaz-Rosshandler, I., Ghazanfar, S., Nichols, J., Wilson, V., Göttgens, B. and Marioni, J. C. (2021). Diverse Routes toward Early Somites in the Mouse Embryo. Dev. Cell 56, 1–13.

Haghverdi, L., Lun, A. T. L., Morgan, M. D. and Marioni, J. C. (2018). Batch effects in single-cell RNA-sequencing data are corrected by matching mutual nearest neighbors. Nat. Biotechnol. 36, 421–427.

Harland, L. T. G., Simon, C. S., Senft, A. D., Costello, I., Greder, L., Imaz-Rosshandler, I., Göttgens, B., Marioni, J. C., Bikoff, E. K., Porcher, C., et al. (2021). The T-box transcription factor Eomesodermin governs haemogenic competence of yolk sac mesodermal progenitors. Nat. Cell Biol. 23, 61–74.

Herrmann, B. G., Labeit, S., Poustka, A., King, T. R. and Lerach, H. (1990). Cloning of the T gene required in mesoderm development in the mouse. Nature 343, 617–622.

Imaz-Rosshandler, I., Rode, C., Guibentif, C., Harland, L. T. G., Ton, M.-L. N., Dhapola, P., Keitley, D., Argelaguet, R., Calero-Nieto, F. J., Nichols, J., et al. (2023). Tracking early mammalian organogenesis – prediction and validation of differentiation trajectories at whole organism scale. Development.

Imaz-Rosshandler, I., Rode, C., Guibentif, C., Harland, L. T. G., Ton, M. L. N., Dhapola, P., Keitley, D., Argelaguet, R., Calero-Nieto, F. J., Nichols, J., et al. (2024). Tracking early mammalian organogenesis - prediction and validation of differentiation trajectories at whole organism scale.

Inman, K. E. and Downs, K. M. (2006). Brachyury is required for elongation and vasculogenesis in the murine allantois. Development 133, 2947–2959.

Inman, K. E. and Downs, K. M. (2007). The murine allantois: Emerging paradigms in development of the mammalian umbilical cord and its relation to the fetus. Genes. (United States) 45, 237–258.

Kinder, S. J., Tsang, T. E., Quinlan, G. A., Hadjantonakis, A., Nagy, A. and Tam, P. P. L. (1999). The orderly allocation of mesodermal cells to the extraembryonic structures and the anteroposterior axis during gastrulation of the mouse embryo. Development 126, 4691–4701.

Koyano-Nakagawa, N. and Garry, D. J. (2017). Etv2 as an essential regulator of mesodermal lineage development. Cardiovasc. Res. 113, 1294–1306.

Koyano-Nakagawa, N., Kweon, J., Iacovino, M., Shi, X., Rasmussen, T. L., Borges, L., Zirbes, K. M., Li, T., Perlingeiro, R. C. R., Kyba, M., et al. (2012). Etv2 is expressed in the yolk sac hematopoietic and endothelial progenitors and regulates Lmo2 gene expression. Stem Cells 30, 1611–1623.

Liu, F., Li, D., Yu, Y. Y. L., Kang, I., Cha, M., Kim, J. Y., Park, C., Watson, D. K., Wang, T. and Choi, K. (2015). Induction of hematopoietic and endothelial cell program orchestrated by ETS transcription factor ER 71/ ETV 2. EMBO Rep. 16, 654–669.

Lugus, J. J., Park, C., Ma, Y. D. and Choi, K. (2009). Both primitive and definitive blood cells are derived from Flk-1 + mesoderm. Blood 113, 563–566.

Lun, A. T. L., McCarthy, D. J. and Marioni, J. C. (2016). A step-by-step workflow for low-level analysis of single-cell RNA-seq data with Bioconductor. F1000Research 5, 2122.

McCarthy, D. J., Campbell, K. R., Lun, A. T. L. and Wills, Q. F. (2017). Scater: Pre-processing, quality control, normalization and visualization of single-cell RNA-seq data in R. Bioinformatics 33, 1179–1186.

Naiche, L. A., Arora, R., Kania, A., Lewandoski, M. and Papaioannou, V. E. (2011). Identity and fate of Tbx4-expressing cells reveal developmental cell fate decisions in the allantois, limb, and external genitalia. Dev. Dyn. 240, 2290–2300.

Nowotschin, S., Costello, I., Piliszek, A., Kwon, G. S., Mao, C. an, Klein, W. H., Robertson, E. J. and Hadjantonakis, A. K. (2013). The T-box transcription factor eomesodermin is essential for AVE induction in the mouse embryo. Genes Dev. 27, 997–1002.

Pijuan-Sala, B., Griffiths, J. A., Guibentif, C., Hiscock, T. W., Jawaid, W., Calero-Nieto, F. J., Mulas, C., Ibarra-Soria, X., Tyser, R. C. V., Ho, D. L. L., et al. (2019). A single-cell molecular map of mouse gastrulation and early organogenesis. Nature 566, 490–495.

Rashbass, P., Cooke, L. A., Herrmann, B. G. and Beddington, R. S. P. (1991). A cell autonomous function of Brachyury in T/T embryonic stem cell chimaeras. Nature 353, 348–351.

Rasmussen, T. L., Kweon, J., Diekmann, M. A., Belema-Bedada, F., Song, Q., Bowlin, K., Shi, X., Ferdous, A., Li, T., Kyba, M., et al. (2011). ER71 directs mesodermal fate decisions during embryogenesis. Development 138, 4801–4812.

Rivera-Pérez, J. A. and Magnuson, T. (2005). Primitive streak formation in mice is preceded by localized activation of Brachyury and Wnt3. Dev. Biol. 288, 363–371.

Schüle, K. M., Weckerle, J., Probst, S., Wehmeyer, A. E., Zissel, L., Schröder, C. M., Tekman, M., Kim, G.-J., Schlägl, I.-M., Sagar, et al. (2023). Eomes restricts Brachyury functions at the onset of mouse gastrulation. Dev. Cell 1627–1642.

Scotti, M., Kherdjemil, Y., Roux, M. and Kmita, M. (2015). A Hoxa13:Cre mouse strain for conditional gene manipulation in developing limb, hindgut, and urogenital system. Genesis 53, 366–376.

Street, K., Risso, D., Fletcher, R. B., Das, D., Ngai, J., Yosef, N., Purdom, E. and Dudoit (2018). Slingshot: cell lineage and pseudotime inference for single-cell transcriptomics. BMC Genomics 19, 1–16.

Tosic, J., Kim, G. J., Pavlovic, M., Schröder, C. M., Mersiowsky, S. L., Barg, M., Hofherr, A., Probst, S., Köttgen, M., Hein, L., et al. (2019). Eomes and Brachyury control pluripotency exit and germ-layer segregation by changing the chromatin state. Nat. Cell Biol. 2019 2112 21, 1518–1531.

Van den Berge, K., Roux de Bézieux, H., Street, K., Saelens, W., Cannoodt, R., Saeys, Y., Dudoit, S. and Clement, L. (2020). Trajectory-based differential expression analysis for single-cell sequencing data. Nat. Commun. 2020 111 11, 1–13.

Wilson, V., Rashbass, P. and Beddington, R. S. P. (1993). Chimeric analysis of T (Brachyury) gene function. Development 117, 1321–1331.

Ying, Q. L., Wray, J., Nichols, J., Batlle-Morera, L., Doble, B., Woodgett, J., Cohen, P. and Smith, A. (2008). The ground state of embryonic stem cell self-renewal. Nature 453, 519–523.

Yzaguirre, A. D.., de Bruijn, M. F. T. R.. and Speck, N. A. (2017). The role of Runx1 in embryonic blood cell formation. Adv. Exp. Med. Biol. 962, 47–64.

Zhao, H. and Choi, K. (2017). A CRISPR screen identifies genes controlling Etv2 threshold expression in murine hemangiogenic fate commitment. Nat. Commun. 2017 81 8, 1–12.

