## Supplementary material for "Eomes directs the formation of spatially and functionally diverse extra-embryonic hematovascular tissues": Table 1

| **Reagent or Resource** | **Source** | **Identifier** | **Usage** | **Dilution** |
| --- | --- | --- | --- | --- |
| **Antibodies** |  |  |  |  |
| Pecam/CD-31 rat IgG antibody | BD Pharmingen | Cat no. 550274, Lot no. 3338195; RRID: AB_393571 | IF |  |
| RFP rabbit IgG | Rockland antibodies | Cat no. 600-401-379, Lot no. 48710; RRID: ABB_2209751 | IF |  |
| Alexa Fluor 594 donkey anti-rabbit IgG | Invitrogen/Life Technology | Cat no. A21207, Lot no. 2313074; RRID: AB_141637 | IF |  |
| Alexa Fluor 488 donkey anti-rat IgG | Invitrogen/Life Technology | Cat no. A21208, Lot no. 2482958; RRID: AB_2535794 | IF |  |
| Anti-T goat polyclonal | Santa Cruz | Cat. no. sc-17745, lot no. A1613; RRID: AB_2200243 | Western | 1:500 |
| Anti-Eomes rat monoclonal | Thermo Fisher | Cat. no.: 14-4875-82, lot no.: 2493129; RRID: AB_11042577 | Western | 1:10,000 |
| Anti-goat IgG polyclonal HRP conjugated | abcam | Cat. no. ab6741, lot no. GR3324245-8; RRID: AB_955424 | Western | 1:2,000 |
| Anti-rat IgG polyclonal HRP conjugated | Cell Signaling | Cat. no. 7077S, lot no. 14; RRID: AB_10694715 | Western | 1:2,000 |
| Pe/Cy7 anti-mouse Flk-1 | BioLegend | Cat. no. 136414, lot no. B358320; RRID: AB_2561607 | Flow | 1:40 |
| BV605 rat anti-mouse PdgfRa | BD Biosciences | Cat. no. 740380, lot no. 4078442; RRID: AB_2740111 | Flow | 1:160 |
| APC anti-mouse Dlk1 | R&D Systems | Cat. no. FAB8634A, lot no. AEPA0224011; RRID: AB_2890004 | Flow | 1:100 |
| **Chemical** |  |  |  |  |
| Vectashield mounting medium with DAPI | Vector Laboratories | Cat no. H-1200, Lot no. ZF1203 |  |  |
| DAPI | BD Pharmigen | 564907 | Flow | 1:5,000 |
| **Cell lines** |  |  |  |  |
| tdTomato-wildtype mESCs | Pijuan-Sala et al., 2019 | CCE or 129? |  |  |
| T-KO tdTomato mESCs | Guibentif et al., 2021 |  |  |  |
| Eo-KO tdTomato mESCs | This paper |  |  |  |
| Eo/T-KO tdTomato mESCs | This paper |  |  |  |
| E14 Runx-Venus Eo^deg/deg^ (clone G9) | Bisia et al., 2023 |  |  |  |
| E14 Runx-Venus Eo^deg/deg^ T-null (clones G9/A11, G9/F9) | This paper |  |  |  |
| **Mouse lines** |  |  |  |  |
| C57BL/6 wildtype mice | Charles River | C57BL/6J |  |  |
| **Oligonucleotides** |  |  |  |  |
| Alt-R® CRISPR-Cas9 tracrRNA | IDT DNA |  | Cas9-based targeting | Resuspended per manufacturer’s instructions |
| Alt-R CRISPR-Cas9 crRNA | IDT DNA | Mm.Cas9.T.1.AD [sequence(PAM)]  [GGTGGTCCACTCGGTACTGC(AGG)] | Cas9-based targeting of T locus | Resuspended per manufacturer’s instructions |
| ssODN | IDT DNA | t*c*tcctccaggcccactcgcagttcgcgttcggtggggtctcccttctcgctgcccgcctgcagctcgctctccacggcgctgagcaggtggtccactcgactagttcatcagtactgcagactcttccctgcgctctctgtgcccggcgagctcatcctcccgccaccctctccacctt*c*c | Repair template for T locus targeting and null allele generation | Resuspended per manufacturer’s instructions |
| Primer AB_289 |  | CTCCGCAGAGTGACCCTTTT | For genotyping T-null clones with AB_290 followed by SpeI restriction digest |  |
| Primer AB_290 |  | TACCTGCCGTTCTTGGTCAC | See above |  |
| Alt-R CRISPR-Cas9 crRNA | Harland et al., 2021 | gRNA#2 [sequence(PAM)]  [GGTTTGCAGCAGGCGATTTG(TGG)] | Cas9-based targeting of Eo locus for exon 2-5 deletion | Resuspended per manufacturer’s instructions |
| Alt-R CRISPR-Cas9 crRNA | Harland et al., 2021 | gRNA#3 [sequence(PAM)]  [AGAGGCATCCCGGCACCCTG(AGG)] |  |  |
| ssODN#2 (SphI) | Harland et al., 2021 | C*C*CTCTTAACCTCCCTCCCCATGCCCTAAATAAACTCTATTCTATACTATTCCATCTTGTGGCTGGTCCCTCAGGGCATGCGCCTGCTGCAAACCCAGGAGCCAGCGGGTCACGTAGATCTGCCCTCAAGGGTTCATTCCCAAATTTCCATC*T*C | Repair template for Eo Δ2-5 null allele generation | Resuspended per manufacturer’s instructions |
| ssODN#3 (SpeI) | Harland et al., 2021 | C*C*CTCTTAACCTCCCTCCCCATGCCCTAAATAAACTCTATTCTATACTATTCCATCTTGTGGCTGGTCCCTCAGGACTAGTGCCTGCTGCAAACCCAGGAGCCAGCGGGTCACGTAGATCTGCCCTCAAGGGTTCATTCCCAAATTTCCATC*T*C | Repair template for Eo Δ2-5 null allele generation | Resuspended per manufacturer’s instructions |
